# An FGF7-FGFR2-KLF4 feedback loop sustains anti-inflammatory signaling in epithelial cells

**DOI:** 10.64898/2026.03.20.711763

**Authors:** Luca Ferrarese, Daria Wüst, Liliana Bento Lopes, Ismail Küçükaylak, Sibilla Sander, Joanne Gerber, Jörn Dengjel, Sabine Werner

## Abstract

Chronic or excessive inflammation are hallmarks of many human diseases, but the endogenous factors that limit inflammatory responses are insufficiently characterized. We discovered a broad anti-inflammatory function of fibroblast growth factor 7 (FGF7), which is mediated via epithelial cels. FGF7 suppressed the expression of pro-inflammatory and immunomodulatory genes in cultured human keratinocytes in the absence or presence of pro-inflammatory stimuli and in a psoriasiform inflammation model in mice. Mechanistically, this involves an FGF7-FGF receptor 2 (FGFR2)-MAPK-Krüppel-like factor 4 (KLF4) signaling axis. FGF7 induced significant alterations in the KLF4 interactome in human keratinocytes and suppressed the transcriptional activity of KLF4 at its pro-inflammatory target genes. Concomitantly, expression of *FGFR2* and downstream signaling components was promoted by KLF4, identifying a KLF4-dependent regulatory feedback loop that sustains anti-inflammatory FGF signaling. These results suggest activation of FGF7-FGFR2-KLF4 signaling as a strategy for the treatment of inflammatory diseases involving the skin or other epithelial tissues and highlight the role of epithelial cells in the control of inflammation.

## INTRODUCTION

FGFs are key regulators of embryonic development, tissue homeostasis, and repair, and their dysregulation is involved in the pathogenesis of major human diseases (Ornitz and Itoh, 2015, 2022). They comprise a family of 22 members, of which most signal via activation of one or more of the four tyrosine kinase FGF receptors (FGFR), designated FGFR1-4. FGF binding induces dimerization and trans-phosphorylation in the intracellular domain of FGFRs and activates three major signaling pathways: the mitogen-activated protein kinase (MAPK) pathway, the phosphoinositide 3-kinase (PI3K) pathway, and the phospholipase Cγ (PLCγ) pathway. This promotes cell proliferation, migration and differentiation, depending on the cell type, the receptor, and the activated signaling pathways (Ornitz and Itoh, 2015, 2022). Some FGFs also have strong cytoprotective activities, in particular FGF7. This FGF exclusively binds and activates the 3b splice variant of FGFR2 (FGFR2b), which is mainly expressed by epithelial cells, but not by mesenchymal or immune cells (Ornitz and Itoh, 2015, 2022). The cytoprotective attributes of FGF7 are clinically relevant, because FGF7 enhances epithelial cell survival and mitigates tissue damage in response to radiation, chemotherapy and other cytotoxic insults (Sadeghi *et al*., 2021). Importantly, Palifermin, a recombinant form of FGF7, is approved for the prevention and treatment of severe mucositis in patients with hematopoietic malignancies that undergo radiation and chemotherapy for bone marrow transplantation (Spielberger *et al*., 2004). The cytoprotective effect of FGF7 involves direct suppression of apoptosis and enhanced expression of genes involved in the detoxification of reactive oxygen species (ROS) and consequent reduction of oxidative stress (Braun *et al*., 2004; Finch and Rubin, 2004; Sadeghi *et al*., 2021; Zheng *et al*., 2024). FGF7 also regulates genes involved in cell-cell adhesion, thereby preserving epithelial integrity (Yang *et al*., 2010; Cai *et al*., 2013). The importance of this activity is highlighted by the phenotype of mice with conditional knockout of *Fgfr1* and *Fgfr2* in keratinocytes (K5-R1/R2 mice), which have lost the capacity to respond to FGF7 and the related FGF10 and FGF22. These mice develop various features resembling the inflammatory skin disease atopic dermatitis (AD), including a defective epidermal barrier resulting from reduced expression of tight junction components. This causes strong transepidermal water loss and skin dryness, which is accompanied by keratinocyte hyperproliferation, overexpression of pro-inflammatory cytokines, and progressive accumulation of immune cells in the skin (Yang *et al*., 2010; Seltmann *et al*., 2018; Sulcova *et al*., 2015). Impaired FGFR2 signaling in keratinocytes is most likely also relevant for AD in humans, because FGFR2 mRNA is significantly down-regulated in basal keratinocytes of lesional skin of these patients (Ferrarese *et al*., 2024). Several other pre-clinical and clinical studies also point to anti-inflammatory effects of FGF7 and FGF10 (Li *et al*., 2024; Geervliet *et al*., 2023; Klufa *et al*., 2019; Finch *et al*., 2013; Coutsouvelis *et al*., 2022), which were thought to be a consequence of the enhanced survival of epithelial cells and improved epithelial integrity. However, a direct anti-inflammatory effect of FGF7 or FGF10 on epithelial cells could not be excluded by these studies. This is relevant, because the previously reported suppression of several interferon-stimulated genes (ISGs) in normal and malignantly transformed keratinocytes and in lung epithelial cells by FGF7 and/or FGF10 (Ferrarese *et al*., 2024; Maddaluno *et al*., 2020; Toriseva *et al*., 2012) and the increased expression of these genes in FGFR2-deficient keratinocytes (Ferrarese *et al*., 2024) supports such a mechanism.

To gain a global overview on the role of FGF7 as a direct or indirect regulator of inflammation, we conducted an unbiased transcriptomic analysis of human keratinocytes under homeostatic conditions and in the presence of pro-inflammatory mediators. The results identify FGF7 as a global negative regulator of inflammatory gene expression in keratinocytes. In functional studies, we discovered an FGF7-FGFR2-MAPK-KLF4 signaling axis, which mediates this effect and induces a positive regulatory loop that sustains anti-inflammatory signaling in epithelial cells.

## RESULTS

### FGF7 suppresses the expression of pro-inflammatory genes in keratinocytes

To determine global changes in the keratinocyte transcriptome that are induced by FGF7 in the presence or absence of pro-inflammatory stimuli, confluent, serum-starved human keratinocytes were pre-treated with vehicle or FGF7 for 3 h, followed by washing and stimulation with vehicle or the pro-inflammatory mediators poly(I:C), which mimics double-stranded RNA, or tumor necrosis factor (TNF)α for 3 h (Fig. 1A-F). Immortalized HaCaT keratinocytes were used for this purpose, because they have a similar FGFR expression pattern as primary human keratinocytes and display a similar responsiveness to FGF7 (Ferrarese *et al*., 2024). RNA from the treated cells was subjected to bulk RNA sequencing (RNA-seq).

**Fig. 1:**
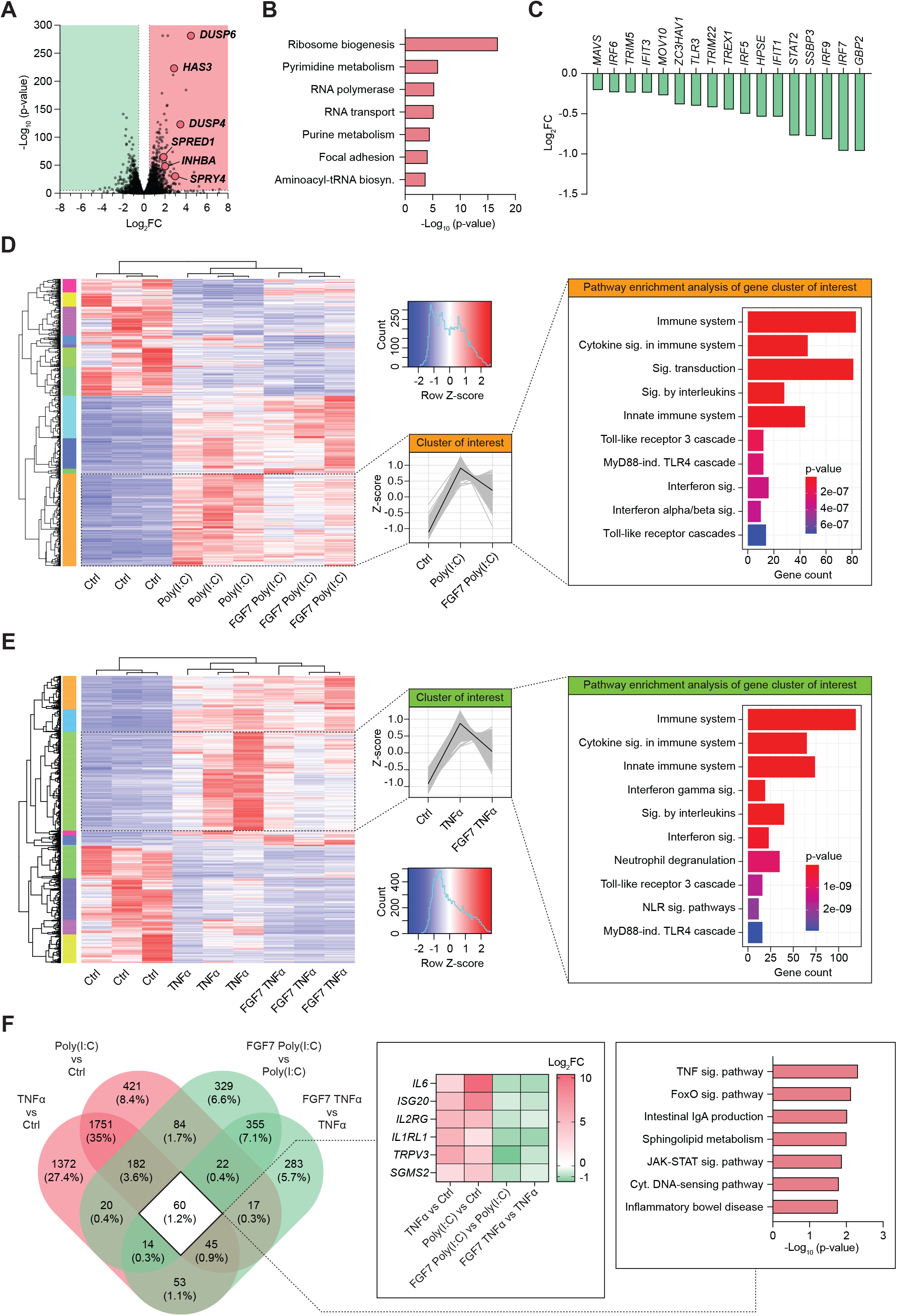
Transcriptome changes of HaCaT keratinocytes in response to FGF7 and pro-inflammatory stimuli. **A)** Volcano blot showing RNA-seq data from serum-starved HaCaT keratinocytes treated with FGF7 or vehicle (PBS) for 3 h. Strongly upregulated genes are indicated (N = 3 per treatment group). **B)** KEGG human pathway enrichment analysis based on the differentially expressed genes (p ≤ 0.05, log2fc ≥ 0.5) in FGF7-*vs*. vehicle-treated HaCaT keratinocytes. **C)** Bar graph showing reduced expression of representative ISGs and pathogen recognition receptors based on RNA-seq data from FGF7-*vs*. vehicle-treated HaCaT keratinocytes. **D, E)** Heatmap illustrating RNA-seq data from serum-starved HaCaT keratinocytes treated with FGF7 or vehicle for 3 h, followed by 3 h treatment with **D)** poly(I:C), **E)** TNFα or vehicle. A cluster of genes with increased expression in response to the respective inflammatory stimulus, but reduced expression upon FGF7 pre-treatment, is highlighted. Right panel shows Reactome pathway enrichment analysis of these gene clusters (N = 3). **F)** Venn diagram showing the number of genes with increased expression in serum-starved HaCaT keratinocytes after 3 h incubation with poly(I:C) or TNFα (red), the number of genes with reduced expression in FGF7-treated cells (green) and the overlap (white), as determined by RNA-seq (N = 3). Cut-off: p ≤ 0.05, log2fc ≥ 0.5 (upregulated) / ≤ −0.5 (suppressed). Heatmap of representative overlapping genes and KEGG human pathway enrichment analysis based on the overlapping genes are shown in the right panel.

In the absence of an inflammatory stimulus, a 3 h treatment with FGF7 affected the expression of 5,790 genes (p ≤ 0.05, false discovery rate (FDR) ≤ 0.05), of which 3,300 exhibited modest effect sizes (log2 fold change (log_2_FC) of ± 0.5). Robust changes were seen in 1,405 upregulated (log_2_FC ≥ 0.5) genes. These include transcripts encoding negative regulators of FGF signaling, such as dual specific phosphatases 4 and 6 (DUSP4, DUSP6), sprouty 4 (SPRY4) and sprouty related EVH1 domain containing 1 (SPRED1) (Ornitz and Itoh, 2022), and the known FGF7 targets hyaluronan synthase 3 (HAS3) (Karvinen *et al*., 2003) and inhibin beta A subunit (INHBA) (Hubner and Werner, 1996) (Fig. 1A). FGF7 also induced the expression of genes encoding tight junction proteins and proteins involved in ROS detoxification (Supplementary Table 1). This is consistent with our previous finding showing the regulation of tight junction genes in mouse epidermis by FGF signaling (Yang *et al*., 2010) and the positive effect of FGF7 on ROS detoxification (Braun *et al*., 2006). Pathway enrichment analysis of the upregulated genes revealed ribosome biogenesis, purine and pyrimidine metabolism, RNA polymerase activity, RNA transport, focal adhesion, and aminoacyl-tRNA biosynthesis as the top hits (Fig. 1B). These findings suggest an overall increase in nucleotide and protein synthesis and cell proliferation upon FGF7 treatment, providing a mechanistic understanding of previous functional data (Rubin *et al*., 1989; Ferrarese *et al*., 2024).

Expression of 1,085 genes decreased in response to FGF7 treatment (log_2_FC ≤ −0.5). Among them are a cluster of genes involved in the interferon response, including *GBP2, IRF5, IRF7, IRF9, IFIT1, IFIT3* and *STAT2*, as well as genes associated with viral infection and pathogen recognition receptor (PRR) signaling (*TRIM22, TLR3* and *TRIM5*) (Fig. 1C).

In response to poly(I:C) or TNFα treatment, we identified prominent clusters of differentially expressed genes (DEGs) (p ≤ 0.05, FDR ≤ 0.05) that were upregulated by the inflammatory stimulus and downregulated by FGF7 (Fig. 1D, E). Pathway enrichment analysis revealed their involvement in immune system processes, cytokine signaling, interleukin signaling, innate immune responses, interferon signaling, and in the toll-like receptor cascade (Fig. 1D, E). Notably, expression of 60 genes was suppressed by FGF7 in both poly(I:C)- and TNFα-treated cells, including *IL6, ISG20, IL2RG, IL1R1, TRPV3* and *SGMS2* (Fig. 1F). Pathway analysis of these overlapping genes highlighted key inflammatory pathways, such as the TNF, the JAK-STAT, and the cytosolic DNA-sensing pathways (Fig. 1F).

These results identify a broad anti-inflammatory activity of FGF7 in keratinocytes under basal conditions and in particular in response to a pro-inflammatory stimulus.

### FGF7 is a negative regulator of interleukin-6 (IL-6) expression in keratinocytes

Given its pivotal role in wound healing and skin inflammation in general (Johnson *et al*., 2020) and its suppression by FGF7 (this study), we focused on IL-6. FGF7 caused a rapid and significant reduction in the basal IL-6 mRNA levels in HaCaT keratinocytes as well as in primary human keratinocytes (HPKs) and in Caco-2 intestinal epithelial cells (Fig. 2A, Supplementary Fig. 1A-C). FGF10, which also activates FGFR2b (Ornitz and Itoh, 2015), induced a comparable decrease (Supplementary Fig. 1A). Pre-treatment with actinomycin D abolished the FGF7 effect on *IL6* and *DUSP6* expression (Fig. 2A, Supplementary Fig. 1D), indicating that it requires transcription. Combined inhibition of MEK and PI3K signaling also prevented FGF7-mediated *IL6* suppression. This is mainly attributed to MEK inhibition, while PI3K inhibition alone had only a minor effect (Fig. 2B, Supplementary Fig. 1E). As expected, FGF7-mediated induction of *DUSP6* expression was MEK-dependent (Li *et al*., 2007), while the induction of *INHBA* expression mainly required PI3K (Supplementary Fig. 1F). Pharmacological inhibition of FGFR kinase activity by BGJ398, AZD4547 and erdafitinib, or genetic knockout of *FGFR2*, promoted *IL6* expression (Fig. 2C, D). This was especially pronounced when *FGFR2* KO HaCaT keratinocytes were treated with TNFα and poly(I:C) (Supplementary Fig. 1G). We also generated HPKs with a CRISPR/Cas9-induced knockout of *FGFR2* and confirmed that FGFR2 deficiency amplifies the increase in *IL6* expression upon treatment with poly(I:C) in primary cells (Supplementary Fig. 1H). FGF7 also attenuated the induction of *IL6* expression in response to the pro-inflammatory stimuli poly(I:C), TNFα, the STING activator 2’3’-cGAMP or the RIG-I agonist 3p-hpRNA (Fig. 2E, F, Supplementary Fig. 1I). Down-regulation of *IL6* expression by FGF7 was equally strong when HaCaT keratinocytes were co-treated for 12 h with FGF7 and TNFα (Supplementary Fig. 1J), demonstrating that FGF7 pre-treatment is not essential. The effect of FGF7 on TNFα-induced IL-6 expression was confirmed at the protein level by ELISA of cell supernatants (Fig. 2G).

**Fig. 2:**
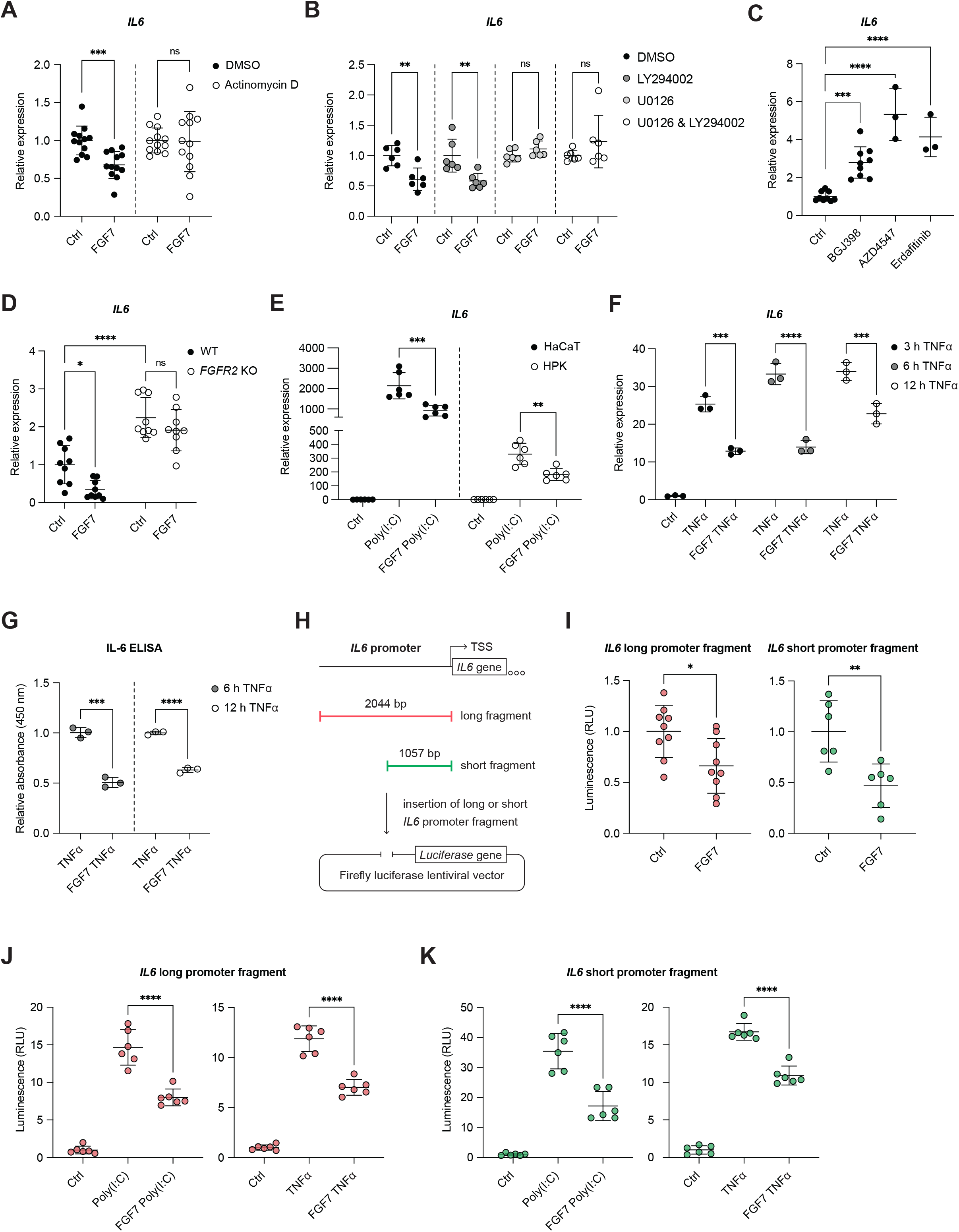
FGF7 suppresses IL-6 expression in keratinocytes. **A)** RT-qPCR for *IL6* relative to *RPL27* using RNA from serum-starved HaCaT keratinocytes, which had been pre-treated for 1 h with actinomycin D or vehicle and incubated for 6 h with FGF7 or vehicle (N = 12 per treatment group). **B)** RT-qPCR for *IL6* using RNA from serum-starved HaCaT keratinocytes, pre-treated for 2 h with LY294002 (PI3K inhibitor), U0126 (MEK1/2 inhibitor) or vehicle, followed by 6 h FGF7 treatment (N = 6). **C)** RT-qPCR for *IL6* using RNA from HaCaT keratinocytes, treated for 6 h with the FGFR kinase inhibitors BGJ398 (N = 9), AZD4547 (N = 3) or erdafitinib (N = 3) in 10% FBS. **D)** RT-qPCR for *IL6* using RNA from serum-starved WT and *FGFR2* KO HaCaT cell lines, treated for 6 h with FGF7 or vehicle (N = 9; 3 different WT and KO cell lines). **E)** RT-qPCR for *IL6* using RNA from serum-starved HaCaT or human primary keratinocytes (HPKs), pre-treated for 3 h with FGF7 or vehicle and incubated for 3 h with poly(I:C) or vehicle (N = 6; HPKs from two donors). **F)** RT-qPCR for *IL6* using RNA from serum-starved HaCaT keratinocytes, pre-treated for 3 h with FGF7 or vehicle and incubated for 3, 6 or 12 h with TNFα or vehicle (N = 3). **G)** Relative IL-6 levels (ELISA) in conditioned medium from serum-starved HaCaT keratinocytes, pre-treated for 3 h with FGF7 or vehicle and incubated for 6 or 12 h with TNFα or vehicle (N = 3). **H)** *IL6* promoter cloning strategy showing 2044 or 1057 base pair (bp) fragments including the transcription start site (TSS), inserted into a firefly luciferase vector. **I)** Luciferase activity in lysates of HaCaT keratinocytes, stably transduced with lentiviruses containing the long or short *IL6* promoter fragment in front of the luciferase gene. Cells were serum-starved and treated for 6 h with FGF7 or vehicle (N = 6). **J, K)** Luciferase activity in lysates of HaCaT keratinocytes, stably transduced with lentiviruses containing the long or short *IL6* promoter fragment in front of the luciferase gene. Cells had been serum-starved and pre-treated for 3 h with FGF7 or vehicle and incubated for 6 h with poly(I:C), TNFα, or vehicle (N = 6). Data information: Graphs show mean and standard deviation (SD). ns: non-significant, *P < 0.05, **P < 0.01, ***P < 0.001, ****P < 0.0001 (Mann-Whitney U test (A, B, I, normalized to respective control), one-way ANOVA with Bonferroni’s multiple comparisons test (C), 2-way ANOVA with Bonferroni’s multiple comparisons test (D; E, normalized to respective control; F; J; K), or Student’s t-test (G, normalized to respective control)).

To verify the transcriptional regulation of *IL6* by FGF7, we performed luciferase reporter assays using a construct in which firefly luciferase expression is driven by the *IL6* promoter. HaCaT keratinocytes were stably transduced with lentiviral vectors containing either a longer (2,044 bp) or shorter (1,057 bp) fragment of the *IL6* promoter (Fig. 2H) in front of the firefly luciferase coding region. The shorter fragment lacks the only classical interferon-stimulated response element (ISRE). FGF7 suppressed luciferase activity under basal conditions and, more notably, in response to inflammatory stimuli (Fig. 2I–K), reinforcing that FGF7 downregulates *IL6* expression at the transcriptional level. The same response was observed with the long and the short reporter construct, demonstrating that the effect of FGF7 is independent of the ISRE.

### FGF7 attenuates pro-inflammatory gene expression in mouse skin

To determine the *in vivo* relevance of our findings, we treated mouse ear skin with Aldara cream, which includes imiquimod and induces a psoriasis-like inflammatory phenotype in mice (van der Fits *et al*., 2009). Concomitantly, we injected FGF7 or vehicle intradermally into the ear skin (Fig. 3A). At day 4 after treatment, early signs of scaling were apparent in Aldara-treated ears (Fig. 3B). FGF7 co-treatment caused a slight increase in ear thickness, whereas transepidermal water loss (TEWL) was not affected (Fig. 3C and D). The increased ear thickness was caused by a significantly thicker epidermis (Fig. 3E, F). Dermal thickness was mildly, but non-significantly reduced (Fig. 3G), suggesting a slight reduction in inflammation-induced swelling. There was a trend toward fewer epidermal CD3^+^ T cells, while the numbers of dermal Ly6G^+^ neutrophils and CD68^+^ macrophages were not altered (Fig. 3H). Despite the increased number of keratinocytes, expression of *Il6, Rsad2* and *Ifnl3* was reduced in six out of seven FGF7-treated ears relative to the PBS-treated ear of the same mouse (Fig. 3I). This effect is most likely underestimated, because we analyzed total skin, while FGF7 only acts on keratinocytes via FGFR2b (Ranieri *et al*., 2018; Werner *et al*., 1993). Together, these results confirm that FGF7 also suppresses inflammatory gene expression in the skin *in vivo*. However, the effect was not sufficient to suppress the immune cell response, possibly because of the down-regulation FGFR2 expression in the inflamed Aldara-treated skin (Fig. 3J).

**Fig. 3:**
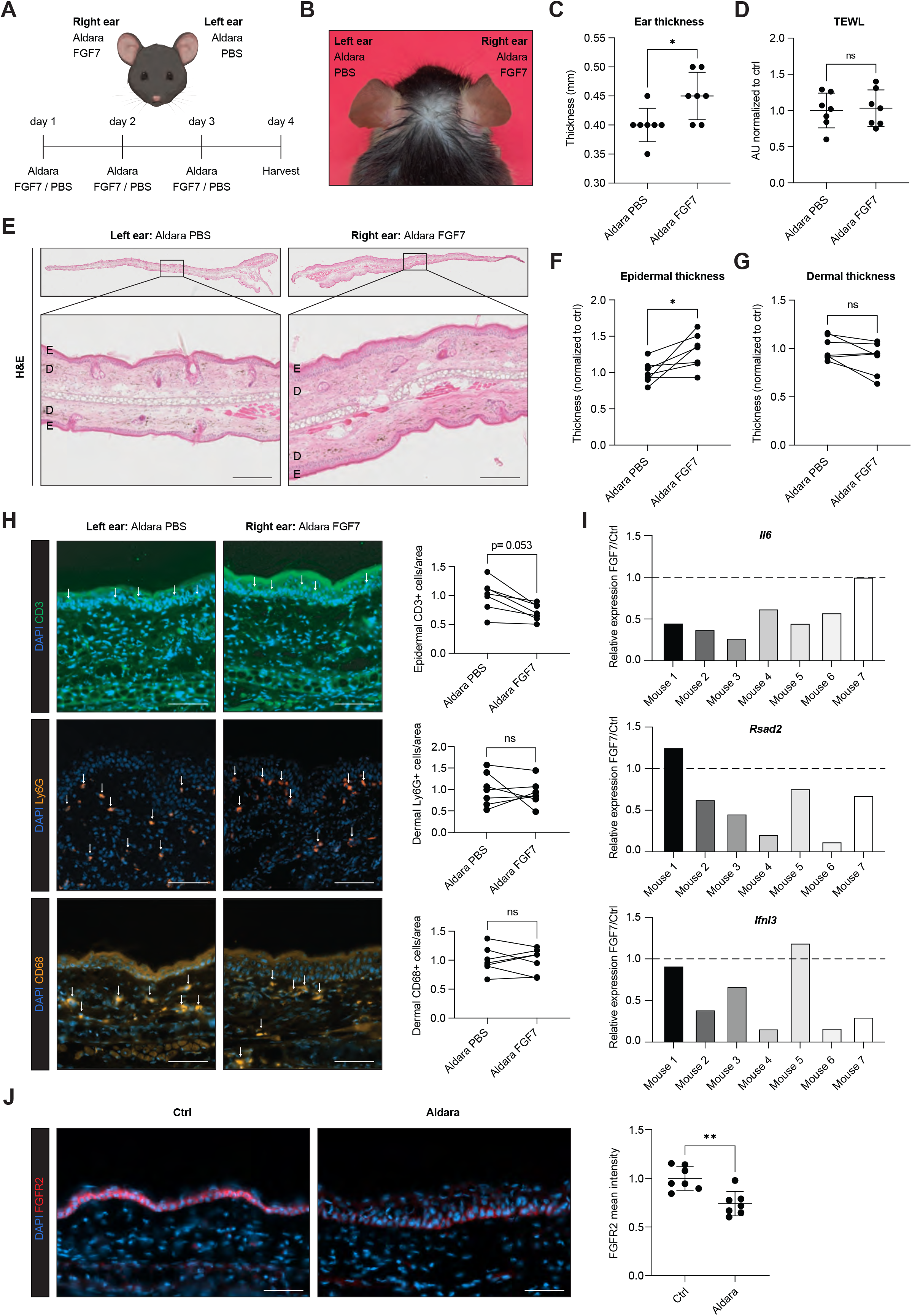
FGF7 suppresses expression of *Il6* and ISGs in a psoriasiform mouse model. **A)** Schematic representation of the experimental set-up, including topical treatment of mouse ear skin with Aldara cream containing 5% imiquimod and intradermal injection of FGF7 or vehicle into the skin of the outer side of the right or left ear, respectively. **B)** Representative picture of dorsal view of a mouse head at day 4, prior to harvest of samples. **C)** Ear thickness (mm) at day 4 (N = 7 mice). **D)** Transepidermal water loss in treated ear skin (TEWL at day 4 (N = 7). **E)** Representative brightfield images of H&E stainings of treated ear skin (N = 7). Scale bars: 150 µm. E: epidermis; D: dermis. **F, G)** Epidermal (F) or dermal (G) thickness based on H&E stainings (E). (N = 7). **H)** Representative immunofluorescence stainings of treated ear skin for CD3 (green), Ly6G and CD68 (orange), counterstained with DAPI (blue). Scale bars: 100 µm. Graphs show quantification of epidermal CD3^+^ cells and dermal Ly6G^+^ and CD68^+^ cells (N = 7). **I)** RT-qPCR for *Il6, Rsad2, Ifnl3* relative to *Rps29* using RNA from the outer part of the treated ear skin. Relative expression in FGF7-treated *vs*. vehicle-treated ear of the same mouse is shown (N = 7). **J)** Representative immunofluorescence images of Aldara- or PBS-treated ears, stained for FGFR2 (red) and counterstained wit DAPI (blue), and quantification of FGFR2 staining intensities (arbitrary units normalized to control). Scale bars: 50 µm. Data information: Graphs show mean and SD. Each line in F-H connects the value of vehicle- and FGF7-treated ears from the same mouse. Bar graphs in (I) depict the FGF7-mediated change in relative expression in Aldara-treated mouse ears compared to the PBS-treated left ear (set to 1). Non-significant (ns), *P < 0.05, **P < 0.01 (Mann-Whitney U test (C; D; F-H, J)).

### FGF7 does not suppress nuclear factor (NF)-κB activity in keratinocytes

Because of the key role of NF-κB in the regulation of pro-inflammatory gene expression (Wullaert *et al*., 2011) and its previously demonstrated regulation by FGF signaling in other cell types (Wang *et al*., 2023; Li *et al*., 2016), we tested if FGF7 affects the activity of this transcription factor (TF). As expected, TNFα promoted phosphorylation of the NF-κB subunit p65 at serine 536, which is required for NF-κB activation (Yang *et al*., 2003; O’Mahony *et al*., 2004), after 10 min and 3 h, and nuclear accumulation of p65 after 30 min, 3 h and 6 h. However, neither p65 phosphorylation nor its nuclear translocation were affected by FGF7 (Supplementary Fig. 2A and B). Furthermore, FGF7 pre-treatment did not alter the DNA-binding activity of NF-κB under homeostatic conditions or after TNFα stimulation (Supplementary Figure 2C). These results suggest that the anti-inflammatory effect of FGF7 in keratinocytes does not involve alterations in NF-κB activity.

### Identification of FGF7-regulated transcription factor motifs

We next applied an unbiased approach to identify the TF that regulate the pro-inflammatory FGF7 target genes. Using Integrated System for Motif Activity Response Analysis (ISMARA), we identified TF motifs that are activated in keratinocytes treated with poly(I:C) or TNFα and exhibit reduced activation upon FGF7 pre-treatment. In the poly(I:C) or TNFα conditions, 92 or 71 TF motifs, respectively, showed increased activity, which was suppressed by FGF7 (z-value ≥⍰±0.5) (Fig. 4A, B). Forty-four TF motifs that were negatively regulated by FGF7 were observed in both inflammatory conditions, including those bound by NF-κB (RELA, REL) and by interferon regulatory factors (IRFs). Notably, motifs for the helicase WRNIP1 and the TFs PATZ1/KLF4 were most strongly affected by FGF7 (Fig. 4C; dark green).

**Fig. 4:**
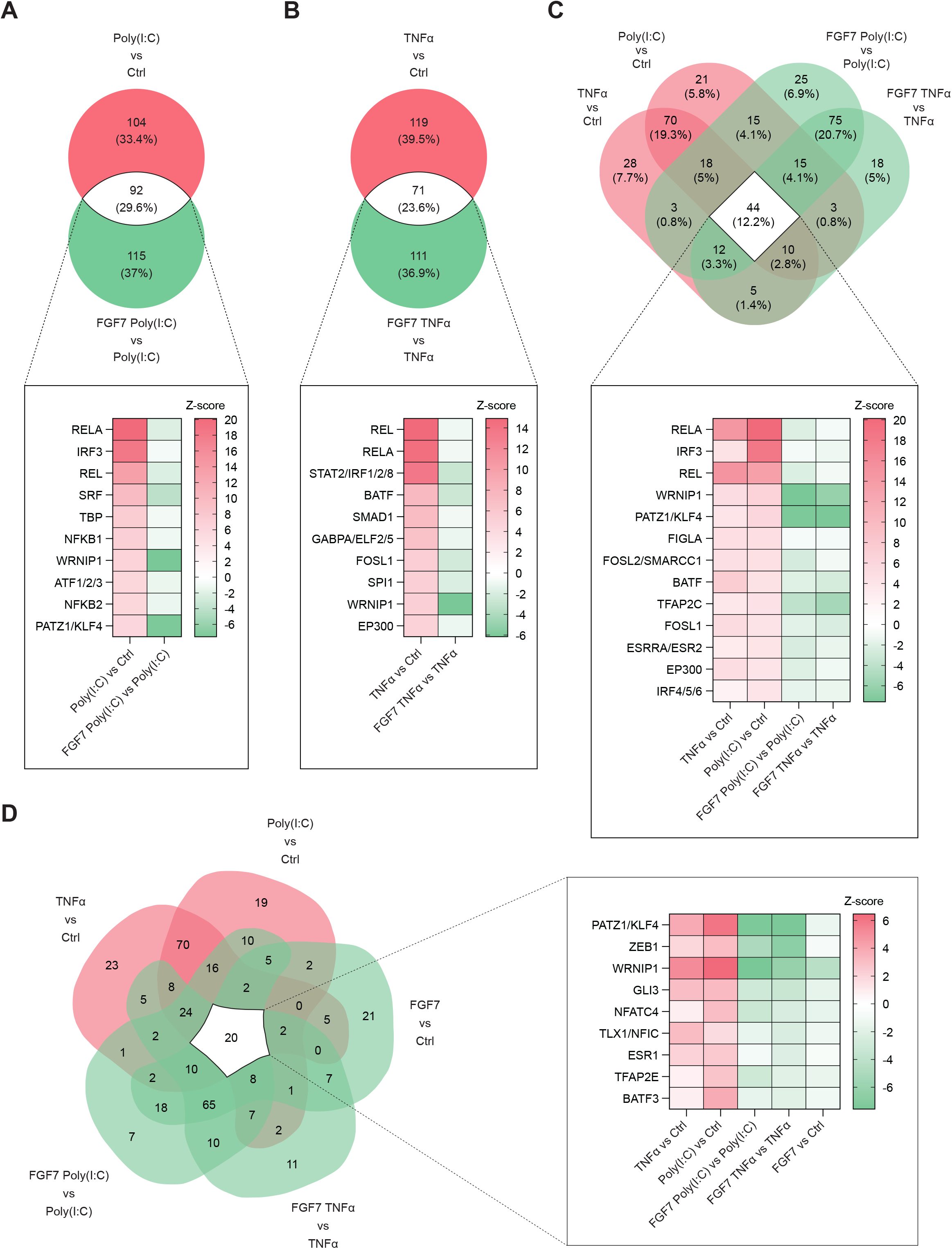
Identification of FGF7-regulated transcription factor (TF) motifs. **A, B)** Venn diagram showing the number of activated TF motifs in serum-starved HaCaT keratinocytes after 3 h incubation with **A)** poly(I:C) or **B)** TNFα (red), the number of TF motifs with reduced activation in FGF7-treated cells (green) and the overlap (white) based on ISMARA. Heat maps show representative overlapping TF motifs. **C, D)** Venn diagrams showing the number of TF motifs that are more (red) or less (green) activated in the different treatment groups based on ISMARA. Heat maps show representative overlapping TF motifs. Cut-off z-value: ≥ +/-0.5. Z-value represents the number of standard deviations by which the motif’s activity deviates from zero. To facilitate interpretation, Z-values of TF motifs exhibiting decreased motif activity were multiplied by −1, resulting in negative values.

Given that FGF7 alone suppresses the expression of ISGs and of *IL6*, we also examined TF motifs that are less activated upon FGF7 treatment in the absence of an inflammatory stimulus. Twenty motifs fulfilled this criterium and showed increased activation in response to poly(I:C) or TNFα, including motifs for PATZ1/KLF4, ZEB1 and WRNIP1 (Fig. 4D). We then analyzed RNA-seq data from HaCaT keratinocytes, which had been treated with FGFR kinase inhibitors (Stefanova *et al*., 2024). ISMARA identified 130 TF motifs, which were activated upon pharmacological FGFR inhibition (z-value ≥⍰0.5) (Supplementary Fig. 3A). Among them were motifs bound by TFs involved in inflammatory responses, such as CEBPB, NF-κB (RELA) and IRFs. Interestingly, TF motifs for IRFs, WRNIP1 and PATZ1/KLF4, which showed reduced activation in all FGF7-treated conditions, exhibited increased activity when FGFR signaling was inhibited (Supplementary Fig. 3A, B).

These computational analyses identified TF motifs that showed increased activation upon loss of FGFR signaling but reduced activation upon FGF7 treatment. The proteins binding to these motifs are potential mediators of the anti-inflammatory effect of FGF7.

### FGF7 suppresses the transcriptional activity of KLF4 at pro-inflammatory target genes

A TF motif that was less activated in response to FGF7 can be bound by the TFs PATZ1 and KLF4. Because of the stronger correlation with KLF4, we focused on this evolutionarily conserved zinc finger-containing TF. KLF4 regulates diverse cellular processes, including proliferation, differentiation and inflammation (Ghaleb and Yang, 2017; Liang *et al*., 2024). It is particularly well known for its role in the induction of pluripotent stem cells (Takahashi and Yamanaka, 2006). KLF4 functions as a transcriptional repressor or activator depending on the context, and its activity is fine-tuned by post-translational modifications and interactions with other proteins (Ghaleb and Yang, 2017). KLF4 mRNA and protein levels were not affected within 1-6 h of FGF7 treatment, and the amounts present in the cytoplasm *vs*. the nucleus were also not obviously altered (Supplementary Fig. 4A-E). However, FGF7 induced significant alterations in the KLF4 interactome. Upon KLF4 pull-down and analysis of interaction partners by mass spectrometry (MS) using a threshold of ≥3 unique tryptic peptides for confident protein identification, we identified a total of 1133 interacting proteins compared to respective negative control IgG pull-downs (Fig. 5A). Among them, 56 proteins were enriched in the interactome upon FGF7 treatment, while 68 proteins were less abundant in the KLF4 interactome of FGF7-treated cells (Fig. 5B). The identified KLF4 binding partners in keratinocytes included proteins that were previously identified in the KLF4 interactome in other human cells according to the BioGRID data base (Oughtred *et al*., 2021) (Supplementary Fig. 5A), demonstrating the validity of the proteomics data. We validated these results in an independent experiment, which largely confirmed the previous data. Importantly, 17 proteins showed a significantly different abundance in the KLF4 interactome upon FGF7 treatment in both experiments (Fig. 5C), again using the threshold of ≥3 unique tryptic peptides. Most of them are associated with transcriptional regulation and/or chromatin remodeling (NF-κB activating protein (NKAP), RuvB-like 2 (RUVBL2), tumor protein 63 (TP63) and syntaxin binding protein 4 (STXBP4), which prevents TP63 degradation (Rokudai *et al*., 2018), zinc finger with KRAB and SCAN domains 1 (ZKSCAN1), AT-rich interaction domain 5B (ARID5B), and ankyrin repeat and KH domain-containing protein 1 (ANKHD1)), or RNA processing (NKAP, cleavage and polyadenylation specificity factor 2 (CPFS1), immunoglobulin μ DNA binding protein 1 (IGHMBP2), up-frameshift protein 1 (UPF1), and the exoribonuclease XRN).

**Fig. 5:**
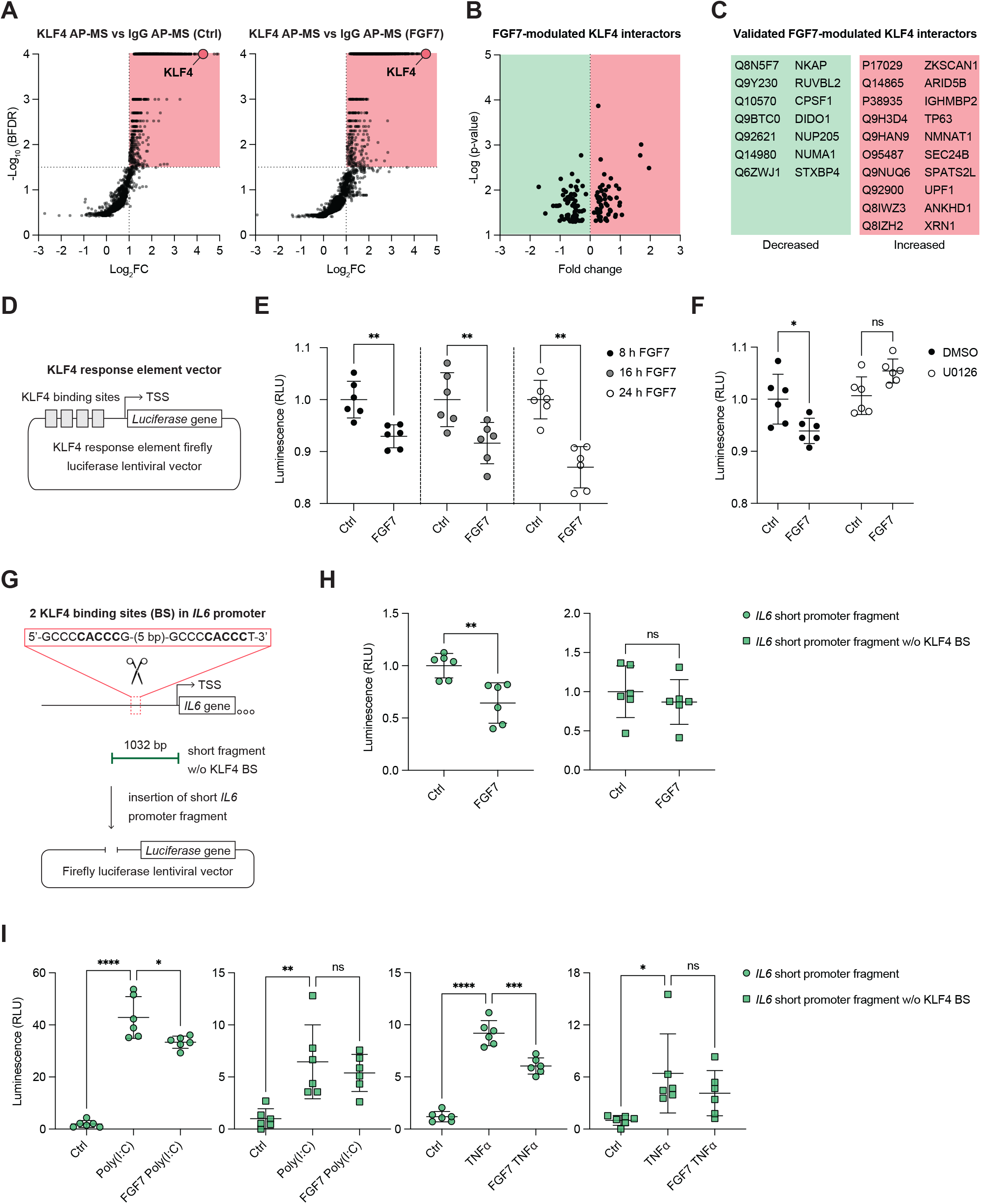
FGF7 alters the KLF4 interactome and suppresses its transcriptional activity. **A)** Volcano plots showing KLF4 affinity purification-mass spectrometry (KLF4 AP-MS) data from serum-starved HaCaT keratinocytes treated with FGF7 for 6 h or vehicle *vs*. the respective IgG control. Enriched prey proteins are located in the upper right quadrant, with KLF4 highlighted (N = 3, one-sided Student’s *t*-test, FDR<0-05). **B)** Volcano plot depicting FGF7-dependent changes in KLF4-associated proteins (KLF4 AP-MS data) under the same experimental conditions. p < 0.05 (two-sided Student’s *t*-test), filtering threshold ≥3 unique tryptic peptides per protein. **C)** Proteins, which also showed a significantly different abundance in the KLF4 interactome upon FGF7 treatment in an independent experiment. **D)** Schematic representation of the luciferase reporter vector harboring 4 KLF4 binding sites in the promoter. **E)** Luciferase activity in lysates of serum-starved HaCaT keratinocytes, stably transduced with lentiviruses including a luciferase reporter gene preceded by KLF4 response elements. Cells were treated for 8, 16 or 24 h with FGF7 or vehicle (N = 6). **F)** Luciferase activity in lysates of HaCaT keratinocytes, stably transduced with lentiviruses containing a luciferase reporter gene preceded by KLF4 response elements. Cells had been serum-starved and pre-treated for 2 h with the MEK1/2 inhibitor U0126 or vehicle, followed by a 6 h treatment with FGF7 or vehicle (N = 6). **G)** *IL6* promoter cloning strategy showing the deletion of two KLF4 binding sites (BS) and the insertion of the *IL6* promoter fragment into the firefly luciferase lentiviral vector. **H)** Luciferase activity in lysates of HaCaT keratinocytes, stably transduced with lentiviruses containing the *IL6* promoter fragment with or without KLF4 BS in front of a luciferase gene. Cells were serum-starved and treated for 6 h with FGF7 or vehicle (N = 6). **I)** Luciferase activity in lysates of serum-starved HaCaT keratinocytes, stably transduced with lentiviruses containing the *IL6* promoter fragment with or without KLF4 BS in front of a luciferase gene. Cells had been serum-starved and pre-treated for 3 h with FGF7 or vehicle and incubated for 6 h with poly(I:C), TNFα, or vehicle (N = 6). Data information: Graphs show mean and SD. Non-significant (ns), *P < 0.05, **P < 0.01; ***P < 0.001; ****P < 0.0001 (Mann-Whitney U test (D, G; normalized to respective control), or 2-way ANOVA with Bonferroni’s multiple comparisons test (E; H).

Overall, these data suggest that FGF7 regulates the transcriptional activity of KLF4, e.g. through altered interaction with other TFs or chromatin remodeling proteins, and possibly also the processing of KLF4-regulated transcripts. To directly test the effect of FGF7 on KLF4’s transcriptional activity, we stably transduced HaCaT keratinocytes with a KLF4 response element-driven firefly luciferase lentiviral vector (Fig. 5D). As expected, luciferase activity was significantly reduced when these cells were transfected with KLF4 siRNA (Supplementary Fig. 5B, C). FGF7 treatment indeed resulted in a significant reduction of luciferase activity after 8, 16 and 24 h (Fig. 5E), which was abolished by inhibition of MEK–ERK signaling (Fig. 5F). Furthermore, deletion of two KLF4 binding sites containing the core KLF4 sequence 5’-CACCC-3’ in the *IL6* promoter construct (Zhang *et al*., 2005) abolished the suppressive effect of FGF7 on luciferase activity (Fig. 5G, H). Most importantly, it also prevented FGF7 from suppressing the *IL6* promoter-driven luciferase activity under inflammatory conditions (Fig. 5I). Together, these findings show that FGF7 suppresses the expression of *IL6* at least in part by regulating the transcriptional activity of KLF4.

### KLF4 is a global regulator of FGF7 target genes

We next scanned the promoter landscape of other FGF7-suppressed genes for KLF4 motifs. Analysis of public ChIP-seq data in the Gene Transcription Regulation Database (GTRD) revealed KLF4 binding sites in the promoters of multiple ISGs that are negatively regulated by FGF7 (Maddaluno *et al*., 2020) (Fig. 6A), which is consistent with the ISMARA data. siRNA-mediated KLF4 knock-down indeed increased the expression of this ISG subset and of *IL6* in HaCaT cells and HPKs (Fig. 6B; Supplementary Fig. 6A-D), phenocopying the transcriptional response observed after genetic or pharmacological inhibition of FGFR signaling (Maddaluno *et al*., 2020; Stefanova *et al*., 2024; Ferrarese *et al*., 2024). KLF4 knock-down also increased the *IL6* promoter-driven luciferase activity, which was abrogated upon deletion of the KLF4 binding sites (Fig. 6C). These results identify KLF4 as a negative regulator of pro-inflammatory gene expression in keratinocytes and as a downstream effector of FGFR signaling.

**Fig. 6:**
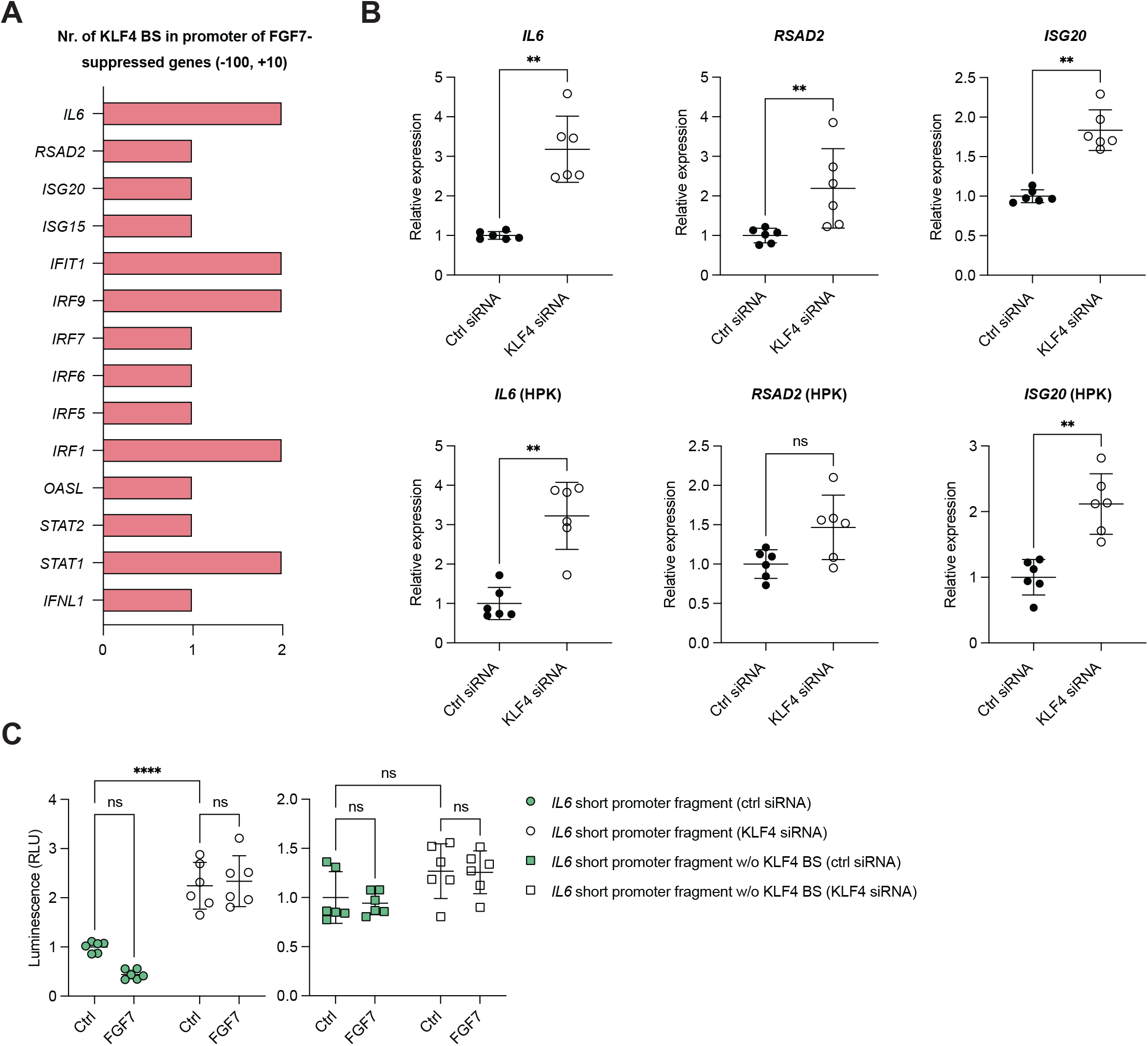
KLF4 directly regulates a cluster of FGF7-suppressed genes. **A)** ChIP-seq data from the Gene Transcription Regulation Database (GTRD) showing the number of KLF4 binding sites (BS) in the promoter regions (−100 to +10 bp relative to the TSS) of FGF7-suppressed genes. **B)** RT-qPCR for *IL6, RSAD2, ISG20* relative to *RPL27* using RNA from serum-starved HaCaT keratinocytes or HPKs, which had been transfected with scrambled (scr) or KLF4 siRNA (N = 6; HPKs from two donors). **C)** Luciferase activity in lysates of HaCaT keratinocytes, stably transduced with lentiviruses containing the *IL6* promoter fragment with or without KLF4 binding sites in front of a luciferase gene. Cells had been transfected with scrambled (scr) or KLF4 siRNA, serum-starved, and treated for 6 h with FGF7 or vehicle (N = 6). Data information: Graphs show mean and SD. Non-significant (ns), **P < 0.01, ****P < 0.0001 (Mann-Whitney U test (B, C)).

### KLF4 is a positive regulator of FGF7-FGFR2 signaling in keratinocytes

Given the importance of regulatory feedback loops in FGFR signaling (Ornitz and Itoh, 2022; Szybowska *et al*., 2021), we determined if KLF4 also regulates the FGF7-FGFR2 pathway itself. Therefore, we monitored FGF7 signaling in HaCaT cells with siRNA-mediated KLF4 knock-down and control cells. Remarkably, KLF4 knock-down reduced the levels of FGFR2 and in particular of its major signaling protein FGFR substrate 2 (FRS2)α (Fig. 7A). This is functionally relevant as shown by the reduced levels of phosphorylated FRS2α and ERK1/2 in FGF7-treated KLF4 knock-down cells (Fig. 7A). Consistently, the FGF7-induced expression of *DUSP6* and *INHBA* was almost abolished (Fig. 7B). While KLF4 knock-down in HaCaT cells and HPKs did not significantly affect FGFR2 mRNA levels at 48 h after siRNA transfection, it reduced the amounts of FRS2α mRNA at this time point (Fig. 7C, D), suggesting that KLF4 may directly regulate *FRS2A* gene expression. We indeed found two KLF4 binding sites within the –100 to +10 bp region relative to the transcription start site and three when expanding the search to –500 to +50 bp. No KLF4 binding sites are present in the *FGFR1* and *FGFR4* gene promoters, one site was identified in the *FGFR3* promoter (–100 to +10 bp), and one in the *FGFR2* promoter (–500 to +50 bp) (Fig. 7E). To determine if KLF4 directly induces the expression of *FRS2A* and *FGFR2*, we delivered a membrane-permeable KLF4-TAT fusion protein into HaCaT keratinocytes. The efficient delivery after 2 h was confirmed by fluorescence staining using a control FITC-TAT protein and by Western blot analysis, which showed the presence of the fusion protein in the cytoplasm and in the nucleus (Supplementary Fig. 7A, B). Eight hours after exposure to KLF4-TAT, FGFR2 and FRS2A mRNA levels were elevated (Fig. 7F). Consistently, basal and FGF7-induced *DUSP6* and *INHBA* expression increased significantly (Supplementary Fig. 7C).

**Fig. 7:**
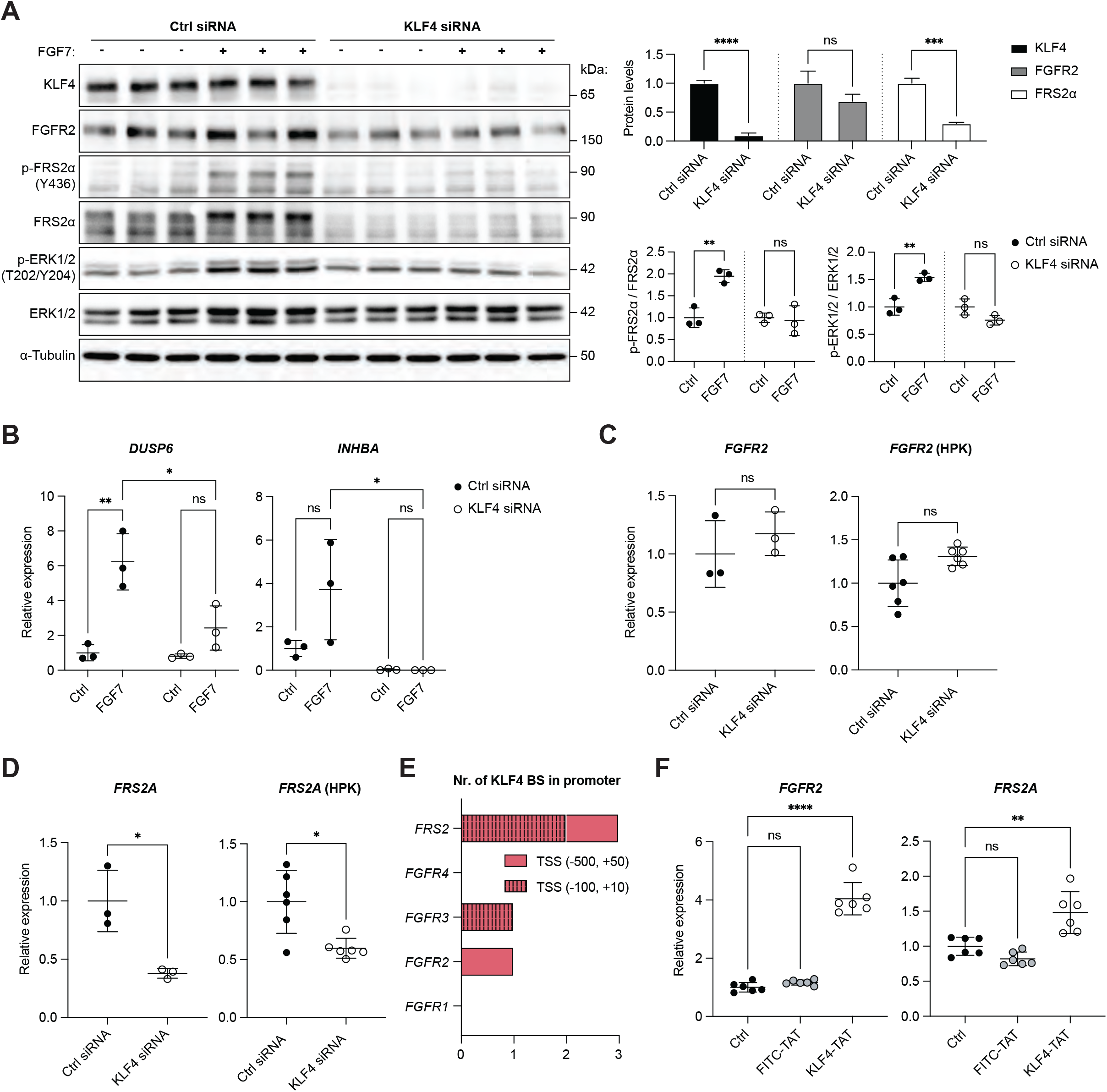
KLF4 directly regulates the FGF7-FGFR2 signaling axis. **A)** Western blot of lysates from serum-starved HaCaT keratinocytes, transfected with scrambled (scr) or KLF4 siRNA mix and treated at 48 h post transfection with FGF7 or vehicle for 15 min. Graphs show densitometric quantification of KLF4, FGFR2 and total FRS2α band intensities normalized to the intensity of α-tubulin (upper panel) or p-FRS2α/FRS2α and p-ERK1/2/total ERK1/2 ratios (lower panels) (N = 3). **B)** RT-qPCR for *DUSP6* and *INHBA* relative to *RPL27* using RNA from serum-starved HaCaT keratinocytes, transfected with scr or KLF4 siRNA and treated at 48 h post transfection with FGF7 or vehicle for 6 h (N = 3). **C, D)** RT-qPCR for *FGFR2* and *FRS2A* using RNA from serum-starved HaCaT keratinocytes or HPKs, transfected with scr or KLF4 siRNA (N = 3-6; HPKs from two donors). **E)** ChIP-seq data from the GTRD showing the number of KLF4 binding sites (BS) in the promoter regions (−500/−100 to +10/+50 bp relative to the TSS) of the *FRS2A* and *FGFR1*-*FGFR4* genes. **F)** RT-qPCR for *FGFR2* and *FRS2A* using RNA from serum-starved HaCaT keratinocytes, incubated with membrane-permeable KLF4-TAT, FITC-TAT, or vehicle for 8 h (N = 6). Data information: Graphs show mean and standard deviation (SD). Non-significant (ns), *P < 0.05, **P < 0.01, ***P < 0.001, ****P < 0.0001 (Student’s t-test (A, normalized to respective control; C, D), one-way ANOVA with Bonferroni’s multiple comparisons test (F) or 2-way ANOVA with Bonferroni’s multiple comparisons test (B)).

Collectively, the loss- and gain-of-function data support a reciprocal circuit in which KLF4 secures the abundance of FGFR2 pathway components (especially FRS2α) to preserve downstream transcriptional outputs, thereby enabling FGF7 to temper epidermal inflammatory programs through FGFR2b-KLF4 signaling (summarized in Fig. 8A). This is of likely relevance *in vivo*, because FGFR2 and KLF4 are co-expressed in basal keratinocytes of the human epidermis (Fig. 8B). Therefore, activation of the FGF7-FGFR2b-KLF4 axis is a promising strategy for the treatment of excessive or chronic skin inflammation.

**Fig. 8:**
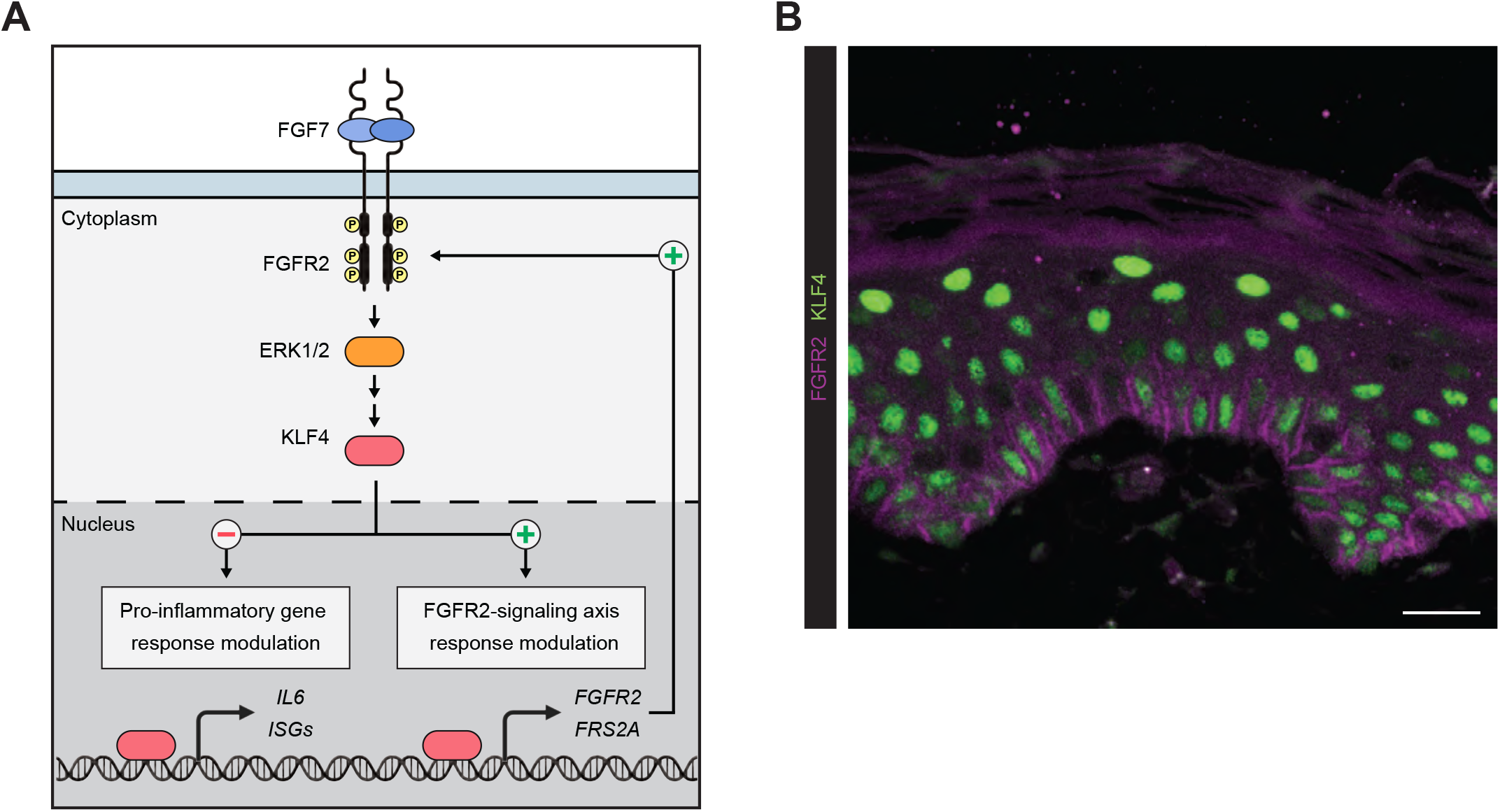
FGF7-FGR2-KLF4 axis in keratinocytes and expression of its components in human skin. **A)** Scheme depicting the FGF7–FGFR2b-ERK1/2-KLF4 signaling axis and its effect on inflammatory gene expression keratinocytes. The icons were obtained from BioRender. Werner, S. (2026) https://BioRender.com/kreppyv. **B)** Representative immunofluorescence stainings of normal human skin for FGFR2 (purple) and KLF4 (green). Scale bar: 20 μm.

## DISCUSSION

We discovered a broad anti-inflammatory and immuno-regulatory effect of FGF7, which involves an FGFR2b-MEK/ERK-KLF4 signaling axis. Our results identify KLF4 as a central component of anti-inflammatory FGF signaling. This finding extends the important role of KLF4 in keratinocytes from the regulation of epidermal barrier function (Segre *et al*., 1999) to direct suppression of pro-inflammatory gene expression in keratinocytes.

A major target of FGF7-KLF4 signaling is *IL6*, which encodes a multifunctional cytokine with a critical role in immune responses and inflammation (Tanaka *et al*., 2014). Elevated levels of IL-6 are implicated in the pathogenesis of various autoimmune and chronic inflammatory diseases (Choy *et al*., 2020; Neurath and Finotto, 2011; Grossman *et al*., 1989; Toshitani *et al*., 1993; Neuner *et al*., 1991; Johnson *et al*., 2020). Our discovery that FGF7 suppresses expression of IL-6 and other pro-inflammatory mediators in keratinocytes helps to understand how growth factor signaling in epithelial cells can modulate inflammatory responses. This is particularly relevant, given that IL-6 not only acts locally, but also exerts systemic pro-inflammatory and immuno-regulatory effects (Rose-John, 2022). However, anti-IL-6 therapy for AD had not only beneficial, but also adverse effects, such as bacterial superinfections (Navarini *et al*., 2011). Therefore, a complete blockade of IL-6 signaling is not recommended. In contrast, FGF7 treatment should only or mainly suppress *IL6* expression in epithelial cells because of the specific expression of its high-affinity receptor FGFR2 in these cells. This could potentially avoid these unwanted side effects, while providing additional cytoprotective effects on keratinocytes. However, down-regulation of FGFR2 in inflamed tissues as seen in lesional skin of AD patients (Ferrarese *et al*., 2024) or in imiquimod-treated mouse skin (this study) may limit the therapeutic effect of FGF7, highlighting the importance of strategies to maintain FGFR2b expression in epithelial cells.

*IL6* expression is also controlled by KLF4 in other cell types, although a positive regulation of this cytokine was observed in most cases (Rosenzweig *et al*., 2013; Xia *et al*., 2021). This suggests a cell type-specific regulation of *IL6* by KLF4 or a specific regulation of KLF4 by FGFR2 signaling. KLF4 also modulates the expression of other genes involved in inflammation, acting as an activator or a repressor depending on the context. In HEK293 cells, KLF4 functions as a repressor by inhibiting the activation of the interferon-β (*IFNb*) promoter and IRF3 recruitment, thereby limiting the production of type I interferons and attenuating antiviral signaling (Luo *et al*., 2016). In the context of sepsis, KLF4 overexpression reduced the expression of TNFα, IL-1β, and IL-6, thereby contributing to a lower inflammatory response and improved survival (Li *et al*., 2021). Our findings suggest that FGF7 reduces the transcriptional activity of KLF4 or promotes its repressor activity, thereby suppressing the expression of *IL6* and most likely other pro-inflammatory genes in keratinocytes. Interestingly, KLF4 cooperated with the glucocorticoid receptor in keratinocytes to induce the expression of TSC223D and ZFP36, which mediate some of the anti-inflammatory effects of glucocorticoids (Sevilla *et al*., 2015). Together, these results point to a global anti-inflammatory effect of KLF4 in keratinocytes by suppressing the expression of pro-inflammatory and promoting the expression of anti-inflammatory genes.

Mechanistically, FGF7 signaling did not alter the amounts of KLF4 in the nucleus. This is different to embryonic stem cells, were KLF4 levels were negatively regulated by phosphorylation through ERK1/2, a key component of canonical FGFR signaling. This led to KLF4’s nuclear export and subsequent proteasomal degradation (Dhaliwal *et al*., 2018; Kim *et al*., 2012). Rather, our proteomics data point to posttranslational regulation of KLF4 by FGF7, which affects its interactome. In the future, it will be important to characterize the FGF7-induced modifications in KLF4 and to identify binding partners at different promoters, which mediate its positive or negative regulation of gene expression. Most of the identified KLF4 interactors identified in this study are involved in transcriptional regulation and chromatin remodeling, suggesting short- and long-term effects of FGF7 on KLF4-mediated transcription. Consistent with this hypothesis, RUVBL2, which showed reduced abundance in the KLF4 interactome of FGF7-treated cells, was previously identified as a co-activator of the related KLF5 transcription factor (Xue *et al*., 2025). In addition to the effect of FGF7 on KLF4-mediated transcription, the altered association of KLF4 with proteins involved in RNA processing in FGF7-treated cells may affect the processing of the KLF4-regulated transcripts.

Our study revealed that FGFR2b and KLF4 act in the same pathway in human keratinocytes to control inflammation. This is of likely relevance *in vivo*, because FGFR2 and KLF4 are co-expressed in normal human skin (this study). *FGFR*2 expression is reduced in basal keratinocytes of lesional skin of AD patients, most likely as a consequence of the overexpression of pro-inflammatory cytokines, which suppressed *FGFR2* expression in cultured keratinocytes (Ferrarese *et al*., 2024). KLF4 levels are also reduced in AD skin based on immunostaining (Sevilla *et al*., 2024). Analysis of our published proteomics data (Koch *et al*., 2023) confirmed this result, and showed a significant down-regulation of KLF4 in the epidermis of lesional *vs*. non-lesional skin of AD patients. This is expected to contribute to the reduced expression of FGFR2 and may even further reduce its signaling capacity as shown by the inhibitory effect of KLF4 knock-down on *FRS2A* expression. However, the activity of KLF4 in AD and other inflammatory skin diseases remains to be determined. This is relevant because FGF7 suppresses *IL6* expression without regulating the abundance or intracellular distribution of KLF4.

In the future, it will be interesting to determine if the FGF7-FGFR2b-KLF4 signaling axis has similar anti-inflammatory effects in other epithelial tissues where FGF7 plays a protective role. This seems likely, because FGF7 also suppressed ISG expression in alveolar epithelial cells (Prince *et al*., 2001), and expression of *IL6* in Caco-2 intestinal epithelial cells (this study), indicating a broad anti-inflammatory effect of FGF7 in different epithelial tissues.

The results obtained in this study enhance our mechanistic understanding of FGF7 activities in epithelial homeostasis and repair: It not only promotes cell proliferation, survival and epidermal barrier function, but also actively suppresses different inflammatory and immuno-regulatory responses in epithelial cells. The identification of FGF7-FGFR2b-KLF4 signaling as a negative regulator of inflammation suggests this axis as a promising target for therapeutic intervention in inflammatory diseases of the skin and other epithelial tissues. Given that FGF7 (Palifermin) is clinically approved for the treatment of mucositis resulting from chemo- and radiation therapy in patients with hematological malignancies (Spielberger *et al*., 2004), an expansion of its applications seems feasible. Furthermore, our results suggest that a combination of Palifermin with approaches that aim to maintain FGFR2 levels in epithelial cells may prolong the activity of Palifermin and could have strong clinical implications.

## MATERIALS AND METHODS

### Cell culture

HaCaT keratinocytes (Boukamp *et al*., 1988), *FGFR2* knockout HaCaT cells (Ferrarese *et al*., 2024), and Caco-2 intestinal epithelial cells were maintained in high-glucose Dulbecco’s Modified Eagle Medium (DMEM, Thermo Fisher Scientific, Waltham, MA) supplemented with 10% fetal bovine serum (FBS, Life Technologies, Carlsbad, CA), at 37⍰°C and 5% CO_2_. Routine mycoplasma screening was conducted using the PCR Mycoplasma Test Kit I/C (PromoCell, Heidelberg, Germany), confirming the absence of contamination.

HPKs were isolated from the foreskin of healthy donors without skin abnormalities as previously described (Sollberger *et al*., 2012). Foreskin samples were anonymously collected, with parental written consent, as part of the University of Zurich (UZH) biobank project. Sample collection and use were approved by the local and cantonal Research Ethics Committees in accordance with the Declaration of Helsinki. HPKs were cultured in Keratinocyte Serum-Free Medium (K-SFM) supplemented with epidermal growth factor (EGF) and bovine pituitary extract (BPE, all from Thermo Fisher Scientific) at 37⍰°C and 5% CO_2_, with 2–3 medium changes per week.

### Generation of HPKs with CRISPR/Cas9-induced knockout of *FGFR2*

Cas9:gRNA ribonucleoprotein particles (RNPs) were assembled using the TrueCut™ HiFi Cas9 Protein (12 pmol, A50576 Thermo Fisher Scientific) and either a custom gRNA targeting *FGFR2* (12 pmol, GCAGTAAATACGGGCCCGAC, Thermo Fisher Scientific) or a TrueGuide™ sgRNA negative control, non-targeting 1 (12 pmol, #A35526, Thermo Fisher Scientific). The complexes were formed by incubation for 20 min at room temperature. HPKs (1.5 x 10^5^ cells) were resuspended in Buffer R (Neon™ NxT Resupension R buffer, #A54298-02, Thermo Fisher Scientific), combined with the Cas9:gRNA RNP complexes and electroporated using a single pulse 1700 V for 20 ms in a Neon™ NxT Electroporation System (Thermo Fisher Scientific). Cells were then plated in antibiotic-free medium, which was refreshed after 48 h. All subsequent experiments were conducted one week after electroporation.

### Treatment of cells with FGF7, FGF10, pharmaceutical inhibitors, TAT-fusion proteins or pro-inflammatory mediators

Cells were seeded into 12-well plates for RNA analysis or into 6-well plates for protein analysis. After reaching confluency, they were either starved overnight (o/n) in DMEM without FBS or treated directly without starvation, as indicated for the respective experiments. Cells were treated with human FGF7 or FGF10 (10 ng/ml, both from Peprotech, Rocky Hill, NJ), poly(I:C) (1 µg/ml, InvivoGen, San Diego, CA), TNFα (20 ng/ml, Peprotech), 2’3’-cGAMP (10 µg/ml, InvivoGen), 3p-hpRNA (1 µg/ml, InvivoGen), actinomycin D (1 µg/ml; Sigma-Aldrich, St. Louis, MO), the FGFR kinase inhibitors BGJ398 (3.6 µM), AZD4547 (10 µM) or erdafitinib (5 µM) (all from Selleckchem, Frankfurt, Germany), the MEK1/2 inhibitor U0126 (10 µM; Calbiochem), the PI3K inhibitor LY294002 (5 µM; Calbiochem), FITC-LC-TAT (100 nM, AnaSpec, Fremont, CA) and/or KLF4-TAT (100 nM, Peprotech).

### Cloning of FFLuc reporter constructs with long and short *IL6* promoter fragments

Genomic human DNA for subsequent *IL6* promoter sequence amplification was isolated from HPKs using the QIAamp DNA Mini Kit (Qiagen, Hilden, Germany) according to the manufacturer’s instructions. PCR was performed using forward (5’-AAG GGC ACA GGT CCT TGA TG-3’) and reverse (5’-TTC TCT TTC GTT CCC GGT GG-3’) primers to amplify the long (2044 bp) *IL6* promoter fragment from genomic DNA. The short (1057 bp) *IL6* promoter fragment was amplified with forward (5’-AAG ACT TAC AGG GAG AGG GAG C-3’) and reverse (5’-TTC TCT TTC GTT CCC GGT GG-3’) primers using the long promoter fragment as template. Next, PCR was performed to create Gibson overhangs on the long (2044 bp) and short (1057 bp) *IL6* promoter fragments using forward (long: 5’-GAT ACT AGT AAG GGC ACA GGT CCT TGA TGT-3’; short: 5’-TCG ATA CTA GTA AGA CTT ACA GGG AGA GGG AGC G-3’) and reverse primers (long: 5’-CTA GCT CTA GAT TCT CTT TCG TTC CCG GTG G-3’; short: 5’-CTA GCT CTA GAT TCT CTT TCG TTC CCG GTG G-3’) that match with the BLIV 2.0 lentivector overhangs. Finally, the constitutive MSCV promoter in a BLIV 2.0 lentiviral vector (System Biosciences, Palo Alto, CA) was replaced by long or short *IL6* promoter fragments via Gibson assembly (New England Biolabs, Ipswich, MA) according to the manufacturer’s instructions.

### siRNA transfection protocol

For transfection, cells were grown to 50-60% confluency. siRNAs were pre-diluted in Opti-MEM® (50 µl per well for 12-well plates and 100 µl per well for 6-well plates), achieving a final concentration of 2.5 µM for both scrambled and target siRNAs. Lipofectamine RNAiMAX (Invitrogen, Carlsbad, CA) was diluted in Opti-MEM® using 50 µl Opti-MEM® + 1.5 µl Lipofectamine per well for 12-well plates and 100 µl Opti-MEM® + 9 µl Lipofectamine per well for 6-well plates. The siRNA and Lipofectamine RNAiMAX mixtures were combined in a 1:1 ratio and incubated at room temperature (RT) for 30 min. Meanwhile, cells were washed once with PBS, and complete DMEM was added. The siRNA-Lipofectamine-RNAiMAX mixture was subsequently added to the cells at a 1:10 dilution. Cells were incubated for 48 h at 37 °C, 5% CO_2_.

### KLF4 binding site deletion

To remove the two KLF4 binding sites (5’-GCCCCACCCG-3’ and 5’-GCCCCACCCT-3’) separated by 5 bp and located 75 bp upstream of the TATA box in the human *IL6* gene promoter, PCR was performed using forward (5’-CAC CCT CCA ACA AAG ATT-3’) and reverse (5’-TGA TTG GAA ACC TTA TTA-3’) primers and the short *IL6*-promoter FFLuc construct as template according to the instructions provided with the Q5® Site-Directed Mutagenesis Kit (New England Biolabs).

### Bacterial transformation and selection

Following the manufacturer’s guidelines, 2 µl of the assembled construct were used to transform 50 µl of NEB 5®-alpha chemically competent *E. coli* (New England Biolabs) or One Shot® TOP10 Chemically Competent *E. coli* (Thermo Fisher Scientific). After subjecting the bacteria to heat shock and allowing them to recover in 1 ml of S.O.C. medium (Thermo Fisher Scientific), 20-150 µl of the transformed culture were spread onto pre-warmed LB agar plates containing 100 µg/ml ampicillin (Sigma-Aldrich). The plates were then incubated o/n at 37°C.

### Screening of insert-containing bacterial colonies, plasmid isolation and sequencing

To identify bacterial colonies that contain the inserted fragment, a colony PCR was conducted. Colonies with the inserted fragment were selected and placed into a 15 ml Falcon® round-bottom polypropylene tube (Corning, New York, NY) containing 5-6 ml of LB medium with antibiotics. Cultures were incubated o/n at 37°C in a shaking incubator. Plasmid isolation was performed using the QIAprep Spin Miniprep kit (Qiagen), following the manufacturer’s protocol. Plasmid DNA was sequenced by Microsynth (Balgach, Switzerland).

### Lentivirus production and transduction of HaCaT keratinocytes

HEK 293T cells were seeded at 40% confluency and transfected the next day with target and packaging plasmids using the Xfect™ Transfection Reagent kit (Clontech Laboratories Inc., Mountain View, CA), following the manufacturer’s protocol. Supernatants containing viruses from transfected HEK 293T cells were collected 48 h post-transfection. Dead cells were removed by centrifugation at 500 g for 5 min at room temperature (RT). The supernatants were then filtered through a 0.45 µm polyethersulfone (PESU) membrane filter (Sarstedt, Nürnbrecht, Germany) and either used immediately for transduction or snap-frozen in 1 ml aliquots with liquid nitrogen and stored at −80°C. For transduction, 2 ml of the viral supernatant supplemented with 8 µg/ml Polybrene® (Santa Cruz Biotechnology, Dallas, TX) were added o/n to HaCaT cells seeded at 50% confluency. The following morning, the cells were washed and cultured in fresh DMEM with 10% FBS. Selection with 1 µg/ml puromycin (Sigma-Aldrich) and/or 1 mg/ml G418 (Thermo Fisher Scientific) was started two days after infection. The medium was replaced every 2-3 days until all non-transduced control cells had died.

### RNA isolation and RT-qPCR

RNA was extracted using the Mini Total Tissue RNA kit (IBI Scientific, Dubuque, IO), which included a DNase digestion step, and the extracted RNA was resuspended in 25-30 μl of diethylpyrocarbonate (DEPC)-treated water. One microgram of RNA was reverse transcribed into cDNA using the iScript cDNA Synthesis Kit (Bio-Rad, Hercules, CA), following the manufacturer’s instructions.

To isolate RNA from mouse ear tissue samples, the tissue was homogenized in 1 ml Trizol® (Life Technologies) using an IKA T25 ULTRA-TURRAX Disperser (IKA Labortechnik, Staufen, Germany). The homogenate was transferred to 2 ml Eppendorf® tubes and incubated at RT for 5 min. Next, 200 µl chloroform (Sigma-Aldrich) were added, mixed by inversion, and incubated for 10 min. Samples were centrifuged at 12,000 g for 15 min at 4 °C to separate the phases. The upper aqueous layer (approximately 400-500 µl) was transferred to a new 2 ml Eppendorf® tube, mixed with 500 µl chloroform, and incubated for another 10 min. After centrifugation under the same conditions, the supernatant (approximately 400-500 µl) was transferred to a 1.5 ml Eppendorf® tube, mixed with 500 µl isopropanol and 1 µl GlycoBlue™ co-precipitant (Invitrogen), and stored at −20 °C o/n for precipitation. The following day, samples were centrifuged at 7,500 g for 5 min at 4 °C, and the supernatant was discarded. The RNA pellet was washed twice with 70% ethanol and centrifuged again at 7,500 g for 5 min at 4 °C. After removing the supernatant, residual liquid was evaporated by a brief incubation at 60 °C. The RNA was dissolved in 25 µl DEPC-ddH_2_O, incubated at 60 °C for 5 min, and stored at −80 °C until use or processed immediately for reverse transcription.

For RT-qPCR amplification, the cDNA was diluted 1:10, and 5 μl of cDNA were combined with 5.5 μl of LightCycler SYBR Green (Roche, Rotkreuz, Switzerland) and 0.5 μl of a 10 μM primer mix. RT-qPCR was carried out in 384-well plates, with all samples run in technical duplicates. Relative gene expression levels were determined using the 2^-ΔΔCt^ method (Livak and Schmittgen, 2001), and expression was normalized to the housekeeping genes *RPL27* (for human cells) or *Rps29* (for mouse tissue). Primer sequences are shown in Table 1.

**Table 1.**
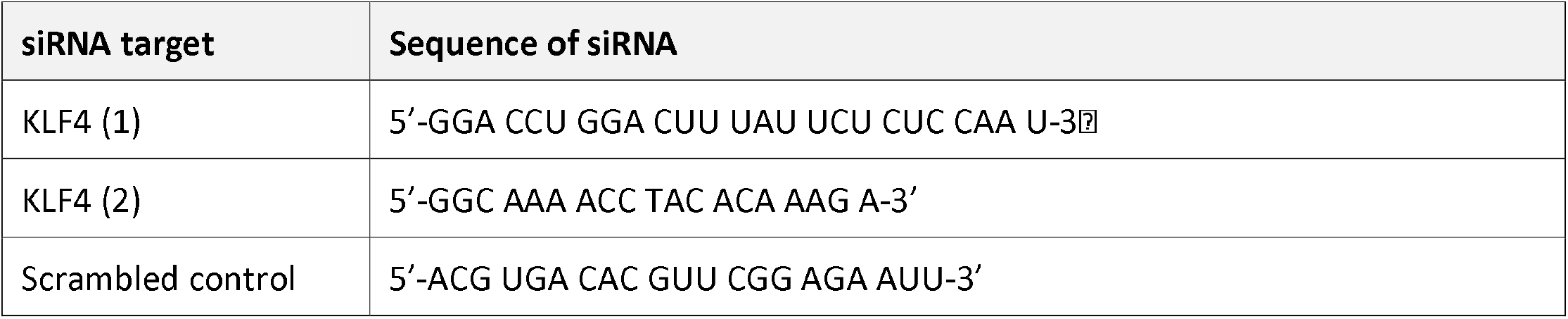
siRNA.

### RNA-seq library preparation and data quantification

RNA was isolated from HaCaT keratinocytes, which had been serum-starved and treated with FGF7 and pro-inflammatory stimuli or respective vehicle using the protocol described above. RNA quality was determined using a Qubit® 1.0 Fluorometer (Life Technologies, Carlsbad, CA) and a Fragment Analyzer (Agilent, Santa Clara, CA). Only samples with a 260/280 nm ratio between 1.8-2.1 and a 28S/18S ratio of 1.5-2 were further processed. The TruSeq Stranded mRNA kit (Illumina, Inc., San Diego, CA) was used for library preparation. Total RNA samples (100-1000 ng) were polyA-enriched and reverse-transcribed into double-stranded cDNA. The cDNA was fragmented, end-repaired, and adenylated before ligation with TruSeq adapters containing unique dual indices (UDI) for multiplexing. Adapter-ligated fragments were selectively enriched by PCR. The quality and quantity of the enriched libraries were validated using the Qubit® Fluorometer and the Fragment Analyzer, yielding a smear with an average fragment size of ~260 bp. Libraries were normalized to 10 nM in Tris-HCl 10 mM, pH 8.5, with 0.1% Tween 20. Cluster generation and sequencing were performed on a NovaSeq6000 System with a single-end 100 bp run configuration, following the NovaSeq workflow and using the NovaSeq6000 Reagent Kit (Illumina, Inc.). RNA-seq data analysis included the following steps: Raw reads were cleaned by removing adapter sequences and poly-x sequences (>9 nt) using fastp (Version 0.20.0) (Chen *et al*., 2018), and reads shorter than 18 nt after trimming were filtered out. High-quality reads were pseudo-aligned to the Human Reference Genome (GRCh38.p13), and gene-level expression was quantified using Kallisto (Version 0.46.1) (Bray *et al*., 2016), with gene models defined by GENCODE release 37. DEGs were identified using the glm approach in DESeq2 (R version 4.1.2, DESeq2 version 1.34.0) (Love *et al*., 2014).

### Dual-luciferase activity assay

Cells stably transduced with lentiviruses encoding Firefly (FFLuc) or Renilla (RLuc) luciferases were grown to full confluency, starved o/n in DMEM, and treated with 10 ng/ml FGF7 or pro-inflammatory mediators for the indicated time periods before assessment of luciferase activity using the Dual-Luciferase® Reporter Assay System (Promega, Madison, WI). After treatment, cells were washed and incubated with 150 µl of Dual-Glo® reagent per well for 30-45 min at 37 °C to lyse the cells. Subsequently, 2 x 50 µl lysate was transferred into each well of a black 96-well plate, and Firefly luciferase luminescence was measured using a GloMax® Discover plate reader (Promega). Following the initial Firefly luminescence measurement, 50 µl of Dual-Glo® Stop & Glo® reagent was added to 50 µl lysate. The plate was incubated for 10 min at RT before measuring Renilla luminescence. The ratio of luminescence from the experimental reporter (FFLuc) to luminescence from the control reporter (RLuc) was calculated to determine the relative luciferase activity. The lentiviral RLuc-mCherry reporter construct was custom-ordered from IDT (Coralville, IA), and the KLF4 response element luciferase reporter construct was purchased from G&P Biosciences (Santa Clara, CA).

### Preparation of protein lysates from cultured cells

Cells were washed twice with ice-cold PBS and lysed with NP-40 lysis buffer (1% NP-40, 150 mM NaCl, 50 mM Tris-HCl), containing PhosSTOP phosphatase inhibitor and cOmplete, EDTA-free protease inhibitor cocktail (both from Roche). Cells were scraped off, and the lysate was placed on a rotating platform for 30 min at 4 °C. Afterwards, samples were centrifuged at 13,800 g for 15 min at 4 °C, and the supernatant was stored on ice or at −20 °C until the protein concentration was measured using bicinchoninic acid (BCA) assay (Thermo Fisher Scientific).

### Western blot

Proteins were separated by SDS-PAGE and transferred onto Protran^™^ 0.2⍰μm nitrocellulose membranes (Amersham, Amersham, UK). After washing with TBS-T (25⍰mM Tris, 137⍰mM NaCl, 2.7⍰mM KCl, 0.1% Tween 20, pH 8.0), unspecific binding sites on the membranes were blocked with 5% bovine serum albumin (BSA; PAN Biotech, Aidenbach, Germany) in TBS-T for 1⍰h at RT, followed by incubation with the primary antibody diluted in TBS-T containing 5% BSA o/n at 4⍰°C. The following day, the membranes were washed three times for 10⍰min each with TBS-T and incubated with the secondary antibody diluted in TBS-T containing 5% BSA for 1⍰h at RT. They were then washed three times for 10⍰min each with TBS-T, and the signal was developed using the Western Bright ECL kit (Advansta, San Jose, CA) and visualized with the FUSION SOLO 6S Western blot imaging system (Vilber Lourmat, Marne-La-Vallé, France). Primary and secondary antibodies are listed in Table 2. Band intensities were quantified using ImageJ/FIJI (National Institutes of Health, Bethesda, MD) and normalized to a housekeeping protein.

**Table 2.**
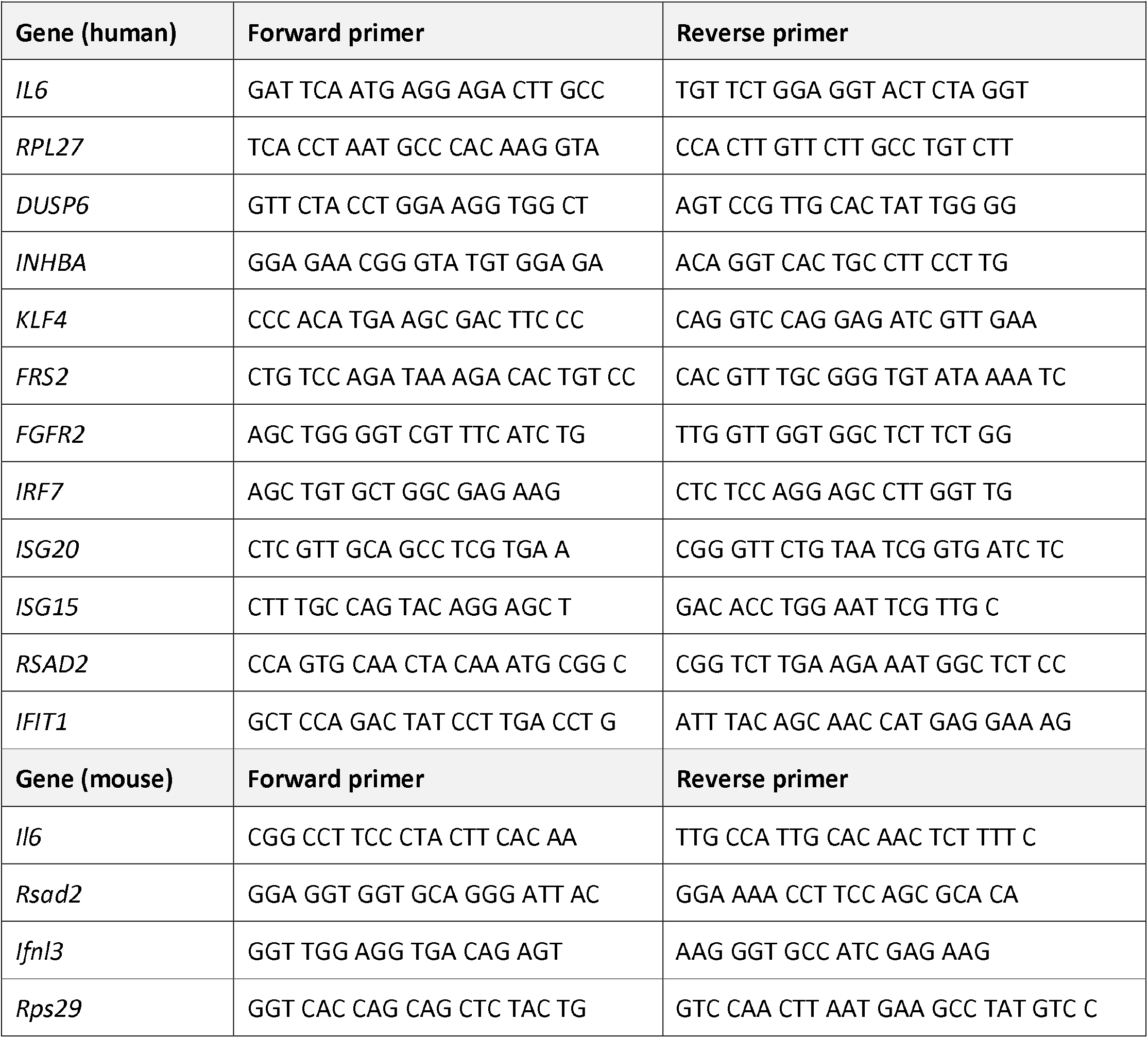
Primer sequences used for RT-qPCR.

### IL-6 ELISA

The IL-6 concentration in conditioned media was measured using the Human IL-6 SimpleStep ELISA Kit (Abcam) according to the manufacturer’s instructions.

### Histological and immunofluorescence stainings

Immunofluorescence staining of cells was performed by seeding cells on glass coverslips in 24-well plates. Samples were washed three times with PBS before being fixed with 4% paraformaldehyde (PFA) for 15 min at RT. Permeabilization and blocking were carried out using Perblock buffer (PBS, 1% BSA, and 0.3% Triton X-100) for 1 h at RT. Primary antibodies (Table 3) were diluted in Perblock buffer. The following day, samples were incubated with the secondary antibody (Table 3) diluted in Perblock buffer for 1 h at RT. Simultaneously, Hoechst (Thermo Fisher Scientific) and phalloidin-iFluor 594 (Abcam) stainings were performed by diluting the stock solutions 1:1000. Samples were washed three times with PBS between each step. Coverslips were mounted on microscopy slides using Mowiol® mounting medium (Sigma-Aldrich) and allowed to air-dry in the dark before imaging with a Zeiss Axiovert Observer.Z1 microscope, equipped with an Axiocam 506 mono camera (both from Carl Zeiss Inc., Oberkochen, Germany).

**Table 3:** Primary and secondary antibodies used for Western blot.

| *Table 3: Primary and secondary antibodies used for Western blot* |  |  |  |
| --- | --- | --- | --- |
| **Antibody** | **Cat. no.** | **Manufacturer** | **Dilution** |
| Goat anti-mouse IgG horseradish peroxidase (HRP) conjugate | W402B | Promega | 1:5000 |
| Goat anti-rabbit IgG HRP conjugate | W4018 | Promega | 1:5000 |
| Mouse anti-glyceraldehyde-3phosphate dehydrogenase (GAPDH) | 5G4 | HyTest, Turku, Finland | 1:1000 |
| Mouse anti-vinculin | V4505 | Sigma-Aldrich | 1:1000 |
| Mouse anti-α-tubulin | T5168 | Sigma-Aldrich | 1:5000 |
| Rabbit anti-histone H3 | ab1791 | Abcam, Cambridge, UK | 1:10000 |
| Rabbit anti-ERK1/2 | 102 | Cell Signaling Technology, Danvers, MA | 1:1000 |
| Rabbit anti-KLF4 | D1F2 | Cell Signaling Technology | 1:1000 |
| Rabbit anti-NF-κB-p65 | 8242 | Cell Signaling Technology | 1:1000 |
| Rabbit anti-FGFR2 | 23328 | Cell Signaling Technology | 1:1000 |
| Rabbit anti-FRS2α | PA5-115253 | Thermo Fisher Scientific | 1:1000 |
| Rabbit anti-AKT | 9272 | Cell Signaling Technology | 1:1000 |
| Rabbit anti-phospho-FRS2α (Tyr436) | 3861 | Cell Signaling Technology | 1:500 |
| Rabbit anti-phospho-ERK1/2 | 9101 | Cell Signaling Technology | 1:1000 |
| Rabbit anti-phospho-NF-κB-p65 | 93H1 | Cell Signaling Technology | 1:1000 |
| Rabbit anti-phospho-AKT (Tyr308) | 9275 | Cell Signaling Technology | 1:1000 |

For immunofluorescence staining of tissue sections, mouse ears were embedded in Tissue-Tek OCT (Sakura Finetek, Torrance, CA) and stored at −80°C until further processing. Cryosections of 10 μm were cut and transferred to SuperFrost Plus slides (Thermo Fisher Scientific). Slides were air-dried for 1 h at RT and subsequently stored at −80°C. They were then immersed in PBS for 10 min, permeabilized with 0.5% Triton X-100 for 20 min and afterwards washed two times for 5 min in PBS. Slides were blocked in blocking buffer (PBS, 10% goat serum, 0.1% Triton X-100, 1% BSA), followed by an overnight incubation with primary antibody in AB buffer (PBS, 0.1% Trition X-100, 1% BSA). After washing three times for 5 min in PBS, slides were incubated with the secondary antibody in AB buffer for 45 min at RT. Primary and secondary antibodies were used according to Table 4. Slides were washed once in PBS, immersed for 5 min in DAPI (Thermo Fisher Scientific) and washed twice in PBS before mounting with Epredia™ Shandon™ Immu-Mount™ (Thermo Fisher Scientific). Images were acquired using the Zeiss Axioscan 7 microscope, equipped with an Axiocam 712 mono camera (both from Carl Zeiss Inc.).

**Table 4:** Primary and secondary antibodies used for immunofluorescence staining.

|  |  |  |  |
| --- | --- | --- | --- |
| **Antibody** | **Cat. no.** | **Manufacturer** | **Dilution** |
| Donkey anti-rabbit IgG AF488 | 71547003 | Jackson ImmunoResearch, West Grove, PA | 1:400 |
| Rabbit anti-NF-κB-p65 | 8242 | Cell Signaling Technology | 1:500 |
| Rabbit anti-CD3 | A0452 | Dako | 1:200 |
| Rat anti-Ly6G | OASB00683 | Aviva Systems Biology, San Diego, CA | 1:200 |
| Rat anti-CD68 | MCA1957 | BioRad | 1:100 |
| Goat anti-rat Cy3 | 112165167 | Jackson ImmunoResearch | 1:500 |

Tissue sections were fixed in ice-cold acetone and stained with hematoxylin (3 min) followed by eosin Y (1 min; Thermo Fisher Scientific). After staining, they were air-dried, mounted with Eukitt® mounting medium (Sigma-Aldrich), and dried o/n at room temperature. Images were acquired using the Zeiss Axioscan 7 microscope, equipped with an Axiocam 712 mono camera (both from Carl Zeiss Inc.).

### Transcription factor binding site prediction

ISMARA (Balwierz *et al*., 2014) was used to predict transcription factor binding sites and to analyze promoter sequences of genes regulated by FGF signaling in the context of inflammation.

### KLF4 antibody conjugation to magnetic beads

To prepare antibody-conjugated beads, 100 µl of magnetic Dynabeads™ Protein A (Invitrogen) were pelleted using a DynaMag™-2 Magnet (Invitrogen) and washed twice with 500 µl of AP binding buffer (0.1 M sodium phosphate buffer, pH 8.0). The beads were resuspended in 80 µl binding buffer, to which 9.21 µg of KLF4 antibody (Cell Signaling) or 9.21 µg of rabbit control IgG antibody (Merck, Darmstadt, Germany) was added. This mixture was incubated at 4 °C for 30 min with gentle agitation. Following antibody binding, the beads were washed three times with 500 µl binding buffer and then twice with AP cross-linking buffer (0.2 M triethanolamine, pH 8.2). They were resuspended in 1 ml of cross-linking buffer containing 6 mg/ml dimethyl pimelimidate dihydrochloride (Sigma-Aldrich) and incubated at RT for 45 min with agitation. To block any remaining reactive groups, the beads were washed once with AP blocking buffer (0.1 M ethanolamine, pH 8.2) and then incubated in 1 ml of blocking buffer for 30 min at RT with agitation. Afterwards, they were washed three times with PBS. Non-cross-linked antibodies were eluted by incubating the beads for 1 min in 1 ml of AP elution buffer (0.1 M glycine-HCl, pH 2.5). Finally, the beads were washed three times with PBS, resuspended in 100 µl of AP lysis buffer (150 mM NaCl, 0.5% NP-40, 10% glycerol, 5 mM HEPES, 5 mM EDTA, pH 7.5) and stored at −20 °C for up to 24 h.

### Preparation of protein lysates for KLF4 affinity purification (AP) mass spectrometry (MS)

O/n starved, FGF7- or vehicle-treated cells at confluency in 15 cm dishes were washed with cold PBS and then lysed in AP lysis buffer. The lysates were sonicated on ice with 10 cycles of 30 s each, followed by 30 s of rest. The samples were then centrifuged at 17,000 g for 10 min at 4 °C, and the supernatant was collected for downstream processing.

Duplicates of each sample, each containing 1 mg of protein, were prepared. To each sample, 10 µl of magnetic Dynabeads™ Protein A coupled to either rabbit control IgG or KLF4 antibody were added. The samples were then adjusted to a final volume of 500 µl with lysis buffer containing protease and phosphatase inhibitors and incubated o/n at 4 °C with agitation. The next day, the beads were pelleted using a DynaMagTM-2 magnet, and the supernatant was collected as the unbound fraction. The beads were washed four times with 800 µl AP lysis buffer without inhibitors and four times with 500 µl AP lysis buffer containing protease and phosphatase inhibitors. Finally, they were subjected to on-bead digestion and MS analysis.

### MS-based proteomics

Proteins were on-bead digested using 500 ng of sequencing-grade modified trypsin (Promega), including tris(2-carboxyethyl)phosphine) (TCEP) and 2-chloroacetamide for reduction and alkylation of cysteines, respectively. Liquid chromatography (LC)-MS analysis of the resulting peptide mixture was conducted on an Orbitrap Exploris 480 (Thermo Fisher Scientific) directly coupled to an ACQUITY UPLC M-Class System (Waters, Milford, MA). MS data acquisition was conducted in data-independent acquisition (DIA) mode: DIA scans were acquired in the Orbitrap at 15,000 resolution covering the m/z range from 350 to 1050 in 70 non-overlapping isolation windows. Precursors were quadrupole isolated at 10 m/z and higher-energy collisional dissociation (HCD) fragmented at a normal collision energy (NCE) of 28 (default charge 3).

MS raw files were processed with Spectronaut version 19 performing peak detection, peptide quantification and identification using a Uniprot human database, August 2023. Carbamidomethyl cysteine was set as fixed modification. Oxidation of methionine, acetylation and phosphorylation were set as variable modification. Three missed cleavages were allowed for trypsin/P as enzyme specificity. Based on a forward-reverse database, protein and peptide FDRs were set to 0.01, minimum peptide length was set to seven, and at least one unique peptide had to be identified. Precursor Q-value, Precursor PEP and Protein Q-value cut-offs were set to 0.01.

### Mouse maintenance and experimentation

Mice were maintained in the ETH Zurich EPIC facility and kept under specific pathogen-free conditions in a 12 h dark/light cycle at 21°C–23°C and 40%-60% humidity. They received food and water *ad libitum*. Mouse maintenance and experimentation had been approved by the veterinary authorities of Zurich, Switzerland (Kantonales Veterinäramt Zurich).

### Induction of psoriasiform inflammation in mice

To induce skin inflammation, 8 to 11-week-old female C57BL/6 mice were topically treated once per day with commercially available Imiquimod (IMQ) cream (Aldara 5%; Viatris Pharma GmbH, Steinhausen, Switzerland) on the outer ear skin, as previously described (van der Fits *et al*., 2009). This was combined with intradermal injection of 2.5 µg recombinant FGF7 (PeproTech) (right ear) or vehicle (left ear) into the outer ear skin using a 33-gauge micro-syringe (Hamilton, Reno, NV). Anesthesia was induced by isoflurane inhalation (Rothacher Medical GmbH, Heitenried, Switzerland) via a nose cone. On day 1, the hairs around the ears were shaved off to facilitate the application of the cream. Ear thickness was measured daily using a Mitutoyo 7301 dial gauge (Mitutoyo, Urdorf, Switzerland) before injection and cream application under short isoflurane anesthesia. After sacrifice, the ears were cut off and divided into two halves. One half was used for histological analysis, while the other half was separated into outer and inner skin parts for RNA isolation.

### Measurement of TEWL

TEWL was assessed using a Tewameter® TM 300 (Courage and Khazaka Electronic GmbH, Cologne, Germany) immediately after euthanizing the mice. Fifteen measurements per ear were performed. The average of these measurements was used for subsequent analysis.

### Tissue collection

Left and right ears were dissected from mouse heads and placed flat on the working surface with the inner ear facing downwards. Each ear was then bisected using single edge blades (AccuTec Blades, Inc., Verona, VA). The upper portion was used for histological analysis and embedded in tissue freezing medium (Leica Biosystems, Wetzlar, Germany) in 25 x 20 x 5 mm Tissue-Tek® Cryomolds® (VWR, Radnor, PA). Samples were frozen on a metal block in liquid nitrogen, and 6 µm sections were obtained using a CryoStar NX70 cryostat (Thermo Fisher Scientific). Three tissue sections per ear were mounted onto SuperFrost Plus™ slides (Thermo Fisher Scientific) and stored at −80 °C until further processing. From the lower portion of the ear, the outer ear skin was removed from the cartilage and the underlying inner ear skin using forceps. The separated ear skin parts were snap-frozen for RNA extraction.

### Statistical analysis

Statistical analysis and generation of graphs were performed using GraphPad Prism 10 software (GraphPad Software Inc, San Diego, CA). Statistical tests were applied as specified in the figure legends.

## ACKNOWLEDGEMENTS

We thank Dr. Petra Boukamp, Leibniz Institute for Environmental Research, Düsseldorf, Germany, for HaCaT keratinocytes, Dr. Hans-Dietmar Beer, University of Zurich, Switzerland, for HPKs, Dr. Tugay Karakaya, University of Zurich, Switzerland, for help with the CRISPR/Cas9-induced knockout of *FGFR2* in HPKs, Dr. Maria Domenica Moccia and Falko Noé, Functional Genomics Center Zurich, for RNA-seq and data analysis, and Dr. Tobias Kockmann, Functional Genomics Center Zurich, for performing the first MS experiment. This work was supported by the Swiss National Science Foundation (grant 310030-212212/1 to S.W.) and the canton and University of Fribourg (to J.D.). J.D. and S.W. are members of the SKINTEGRITY.CH collaborative research program.

## DATA AVAILABILITY STATEMENT

All data are shown in the Figures and Supplementary Figures. Raw data will be provided by the corresponding author upon request. Original RNA-seq files were deposited in the Gene Expression Omnibus (GEO) (GSE302370). MS proteomics data were deposited to the ProteomeXchange Consortium via the PRIDE partner repository (Perez-Riverol *et al*., 2025) with the dataset identifier PXD075609.

## AUTHOR CONTRIBUTIONS

Conceptualization: LF, SW; JD; Investigation: LF, DW, LVL, IK, SS, JG; Methodology: LF, DW, SS; Data Curation: LF; Formal Analysis: LF, IK, SS, JD; Funding Acquisition: SW, JD; Resources: SW, JD; Supervision: SW, JD; Visualization: LF; Writing - Original Draft Preparation: LF, SW,

