## Supplementary Figures, Table and Legends for "An FGF7-FGFR2-KLF4 feedback loop sustains anti-inflammatory signaling in epithelial cells"

### SUPPLEMENTARY FIGURE LEGENDS AND SUPPLEMENTARY TABLE

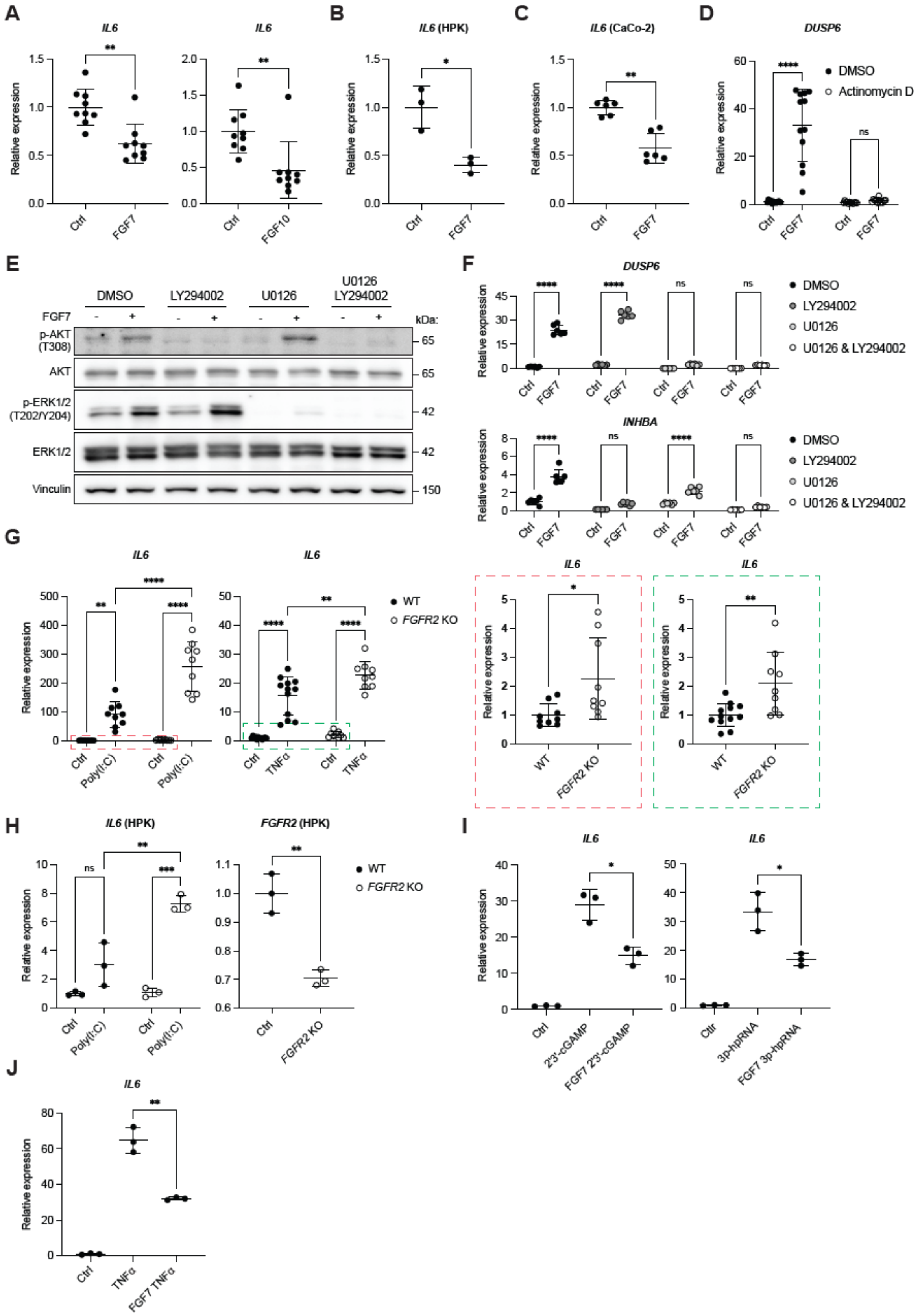

A
IL6
IL6
B
IL6 (HPK)
C
IL6 (CaCo-2)
D
DUSP6
DMSO
Actinomycin D
E
DMSO
LY294002
U0126
U0126 LY294002
FGF7
kDa:
p-AKT (T308)
AKT
p-ERK1/2 (T202/Y204)
ERK1/2
Vinculin
F
DUSP6
INHBA
U0126 & LY294002
G
IL6
WT
FGFR2 KO
H
IL6 (HPK)
FGFR2 (HPK)
I
IL6
2'3'-cGAMP
FGF7 2'3'-cGAMP
3p-hpRNA
FGF7 3p-hpRNA
J
IL6
TNFα
FGF7 TNFα
Relative expression

**Supplementary Fig. 1: FGF7 suppresses *IL6* expression in human keratinocytes**

**A-C)** RT-qPCR for *IL6* relative to *RPL27* using RNA from serum-starved HaCaT keratinocytes (A), human primary keratinocytes (HPKs) (B) or Caco-2 intestinal epithelial cells (C) incubated for 6 h with FGF7, FGF10 or vehicle as indicated (N = 9 per treatment for HaCaT cells; N = 3 for HPKs (single donor), N = 6 for Caco-2 cells).

**D)** RT-qPCR for *DUSP6* using RNA from serum-starved HaCaT keratinocytes, which had been pre-treated for 1 h with actinomycin D or vehicle and incubated for 6 h with FGF7 or vehicle (N = 12).

**E)** Western blot of lysates from starved HaCaT cells, pre-treated for 2 h with LY294002, U0126, a combination of both inhibitors, or vehicle (DMSO), before stimulation with FGF7 or vehicle for 10 min. Membranes were probed with antibodies against total and phosphorylated AKT and ERK1/2 or vinculin (additional loading control).

**F)** RT-qPCR for *IL6* using RNA from serum-starved HaCaT keratinocytes, pre-treated for 2 h with LY294002, U0126, a combination of both inhibitors, or vehicle (DMSO), and incubated for 6 h with FGF7 or vehicle (N = 6).

**G)** RT-qPCR for *IL6* using RNA from WT and *FGFR2* KO HaCaT cell lines, incubated for 6 h with poly(I:C), TNFα, or vehicle (N = 9-12 using 3-4 different WT and KO cell lines). Values of non-treated conditions (red and green dotted boxes) are replotted in the right-hand panels.

**H)** RT-qPCR for *IL6* and *FGFR2* using RNA from WT and *FGFR2* KO HPKs, incubated for 6 h with poly(I:C) or vehicle (N = 3 per genotype and treatment group).

**I)** RT-qPCR for *IL6* using RNA from serum-starved HaCaT keratinocytes, pre-treated for 3 h with FGF7 or vehicle and incubated for 3 h with 2`3`-cGAMP, 3p-hpRNA, or vehicle (N = 3).

**J)** RT-qPCR for *IL6* using RNA from serum-starved HaCaT keratinocytes, co-treated for 12 h with FGF7 and TNFα or vehicle (N = 3).

Data information: Graphs show mean and SD. ns: non-significant, *P < 0.05, **P < 0.01, ***P < 0.001, ****P < 0.0001 (Mann-Whitney U test (A; C), Student`s t-test (B; H, right graph) or 2-way ANOVA with Bonferroni`s multiple comparisons test (D; F; G; H, left graph; J)).

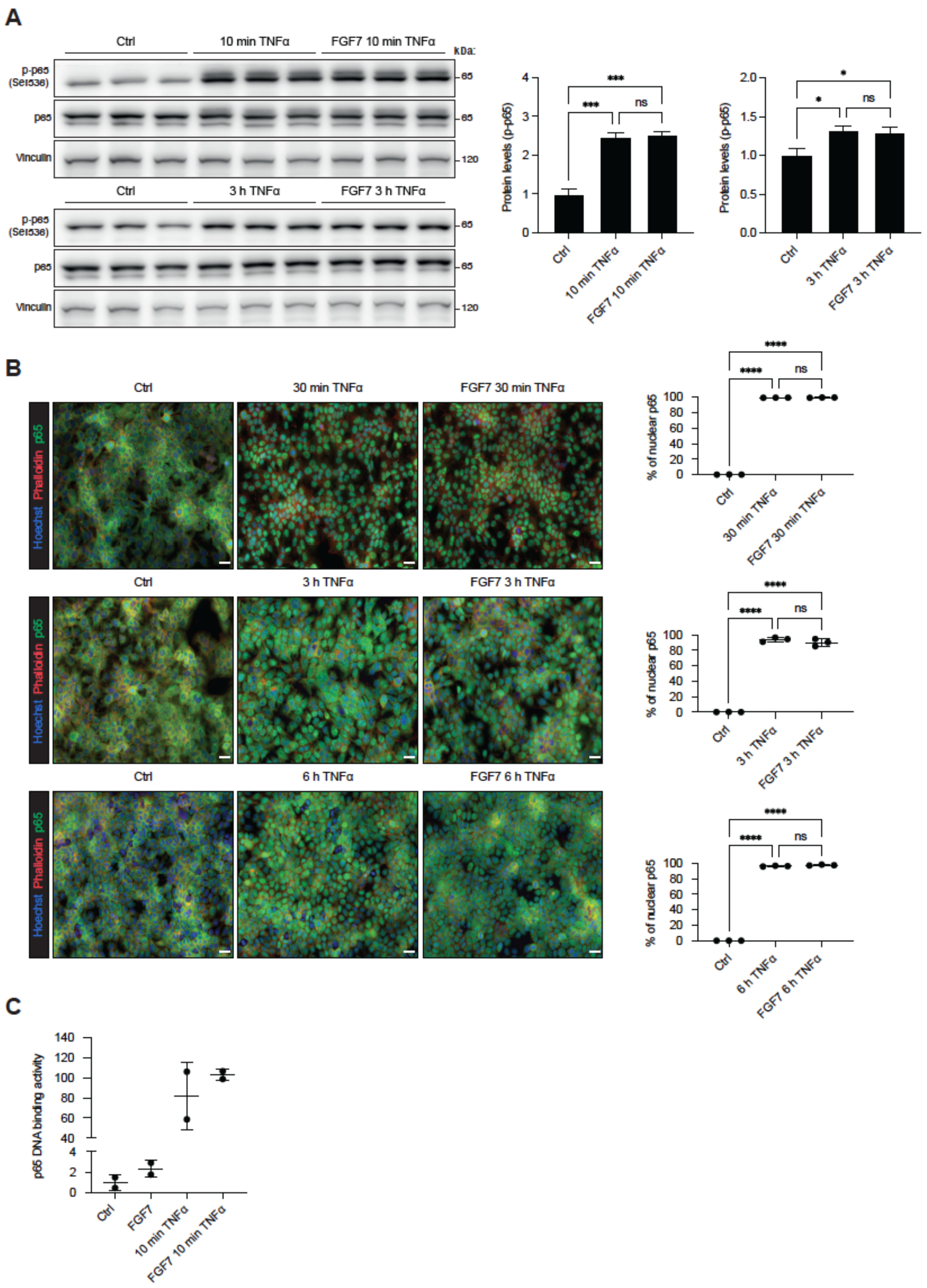


**Supplementary Fig. 2: FGF7 does not affect NF-κB activity in human keratinocytes**

**A)** Western blot of lysates from serum-starved HaCaT keratinocytes, pre-treated for 3 h with FGF7 or vehicle and incubated for 10 min or 3 h with TNFα. Membranes were probed with antibodies against NF-κB-p65 (Ser536), total p65, or vinculin (loading control). Graphs show densitometric quantification of p-p65 relative to total p65 (N = 3).

**B)** Representative IF stainings of serum-starved HaCaT keratinocytes, pre-treated for 3 h with FGF7 or vehicle and incubated for 30 min, 3 h or 6 h with TNFα, stained for p65 (green), and counterstained with phalloidin-iFluor 594 (red) and Hoechst (blue). Scale bars: 50 µm. The percentage of p65-positive nuclei is shown in the graphs (N = 3).
**C)** NF-κB-p65 DNA binding activity in nuclear lysates from starved HaCaT keratinocytes, pre-treated for 3 h with FGF7 or vehicle and incubated for 10 min with TNFα, determined by an ELISA-based assay (N = 2).
Data information: Graphs show mean and SD. Non-significant (ns), $*P < 0.05$, $***P < 0.001$, $****P < 0.0001$ (2-way ANOVA with Bonferroni`s multiple comparisons test).

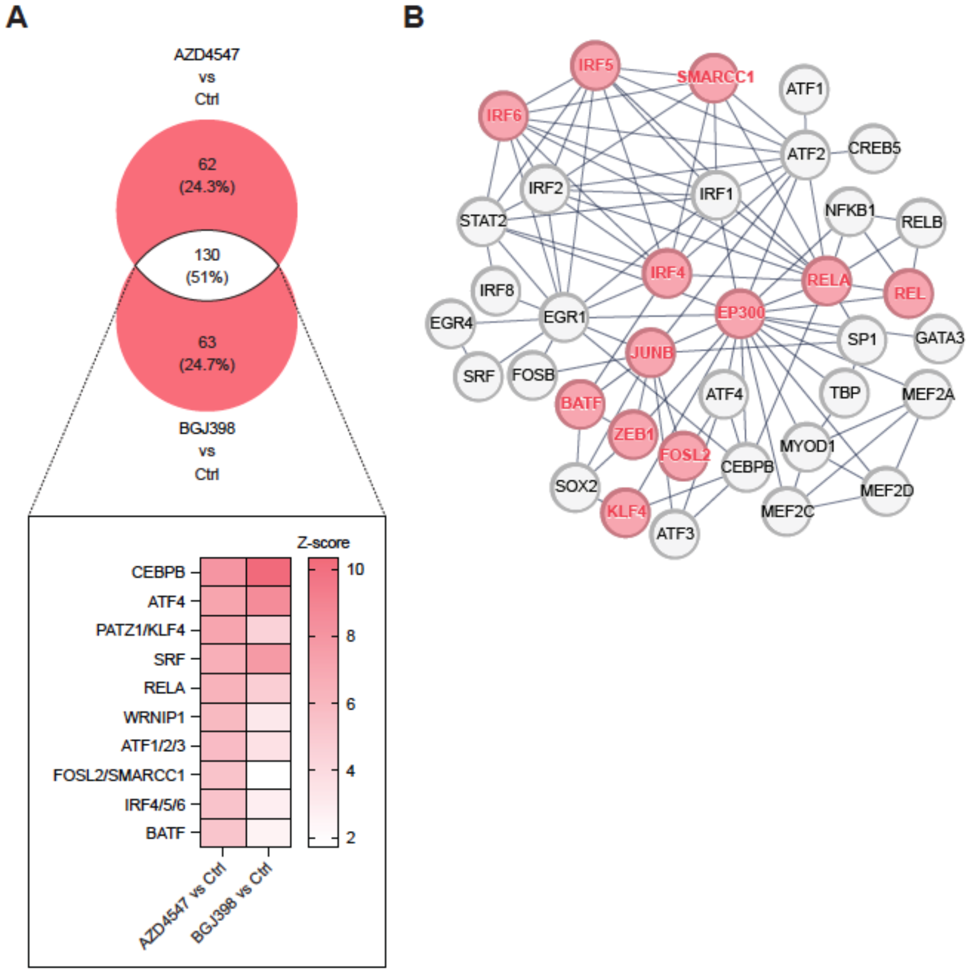


**Supplementary Fig. 3: TF motifs with increased activity upon pharmacological inhibition of FGFR signaling**

**A)** Venn diagram showing the number of activated TF motifs in HaCaT keratinocytes after 5 h treatment with AZD4547 or BGJ398 (red) and overlap (white), based on ISMARA of RNA-seq data (Stefanova *et al.*, 2024). Heat map shows representative overlapping TF motifs.

**B)** STRING analysis showing physical or functional interactions of TFs, which bind to the TF motifs that show increased activation upon pharmacological FGFR inhibition based on ISMARA. TF highlighted in red bind to TF motifs, which show increased activation upon pharmacological FGFR inhibition as well as upon treatment with inflammatory stimuli, while showing reduced activation when pre-treated with FGF7.

Cut-off z-value: ≥ 0.5. Z-value represents the number of standard deviations by which the motif's activity deviates from zero.

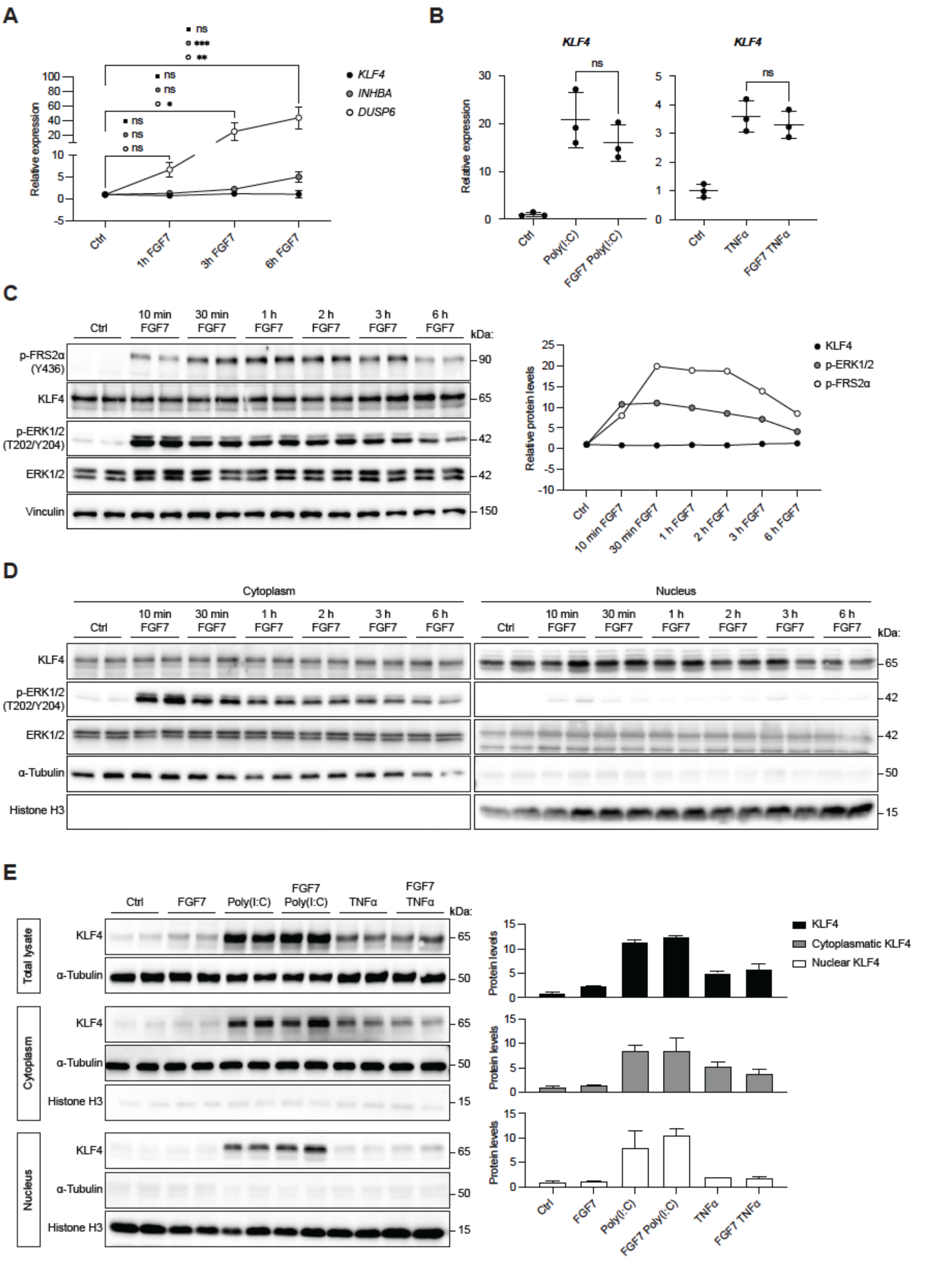


**Supplementary Fig. 4: Short-term FGF7 treatment does not affect KLF4 expression or intracellular localization in HaCaT cells**

**A)** RT-qPCR for *KLF4*, *INHBA* and *DUSP6* relative to *RPL27* using RNA from serum-starved HaCaT keratinocytes, treated for 1 h, 3 h or 6 h with FGF7 or vehicle (N = 3).

**B)** RT-qPCR for *KLF4* using RNA from serum-starved HaCaT keratinocytes, pre-treated for 3 h with FGF7 or vehicle and incubated for 3 h with poly(I:C), TNFα, or vehicle (N = 3).
**C)** Western blot of lysates from serum-starved HaCaT keratinocytes, treated with FGF7 or vehicle for 10 min, 30 min, 1 h, 2 h, 3 h or 6 h. Graph shows average of densitometric quantification of the KLF4/vinculin, p-ERK1/2/total ERK1/2 and p-FRS2α/vinculin ratios (N = 2).
**D)** Western blot of cytoplasmic and nuclear fractions from serum-starved HaCaT keratinocytes, treated with FGF7 or vehicle for 10 min, 30 min, 1 h, 2 h, 3 h or 6 h.
**E)** Western blot of total, cytoplasmic and nuclear lysates of serum-starved HaCaT keratinocytes, pre-treated for 3 h with FGF7 or vehicle and incubated for 3 h with poly(I:C), TNFα or vehicle. Graphs show densitometric quantification of KLF4/α-tubulin ratios in total and cytoplasmic lysates and KLF4/histone H3 ratio in nuclear lysates (N = 2).
Data information: Graphs show mean and SD. Non-significant (ns), *P < 0.05, **P < 0.01, ***P < 0.001 (One-way ANOVA with Bonferroni`s multiple comparisons test (A, normalized to respective control), or 2-way ANOVA with Bonferroni`s multiple comparisons test (B)).

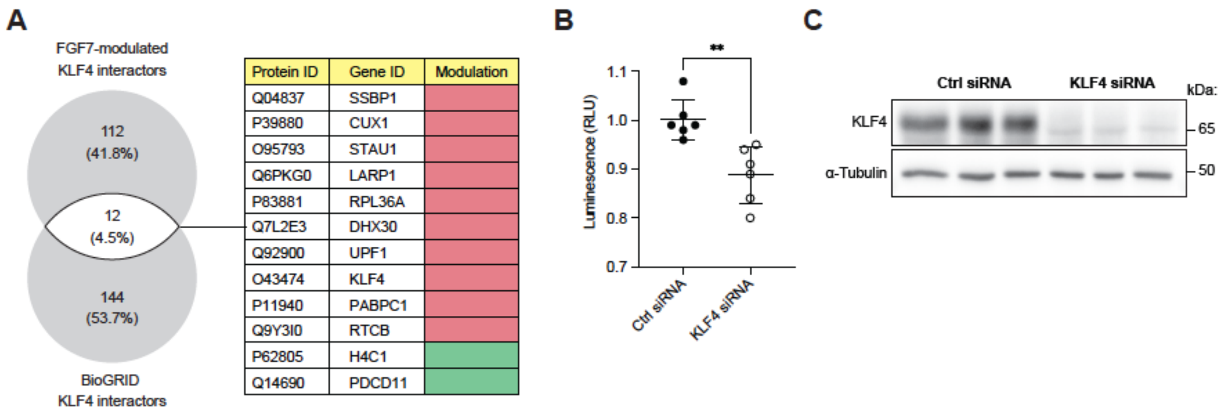


**Supplementary Fig. 5: FGF7 alters the KLF4 interactome**

**A)** Venn diagram illustrating the overlap between FGF7-modulated, KLF4-associated proteins in HaCaT cells and known KLF4 interactors in other human cells curated in the BioGRID database. STRING analysis showing physical or functional interactions of overlapping proteins.

**B)** Luciferase activity in lysates of HaCaT keratinocytes, stably transduced with lentiviruses containing KLF4 response elements in front of the luciferase gene and transfected with scrambled (scr) or KLF4 siRNA (N = 6).

**C)** Western blot of lysates of HaCaT keratinocytes, transfected with scrambled (scr) or KLF4 siRNA.

Data information: Graph shows mean and SD. **P < 0.01 (Mann-Whitney U test (B).

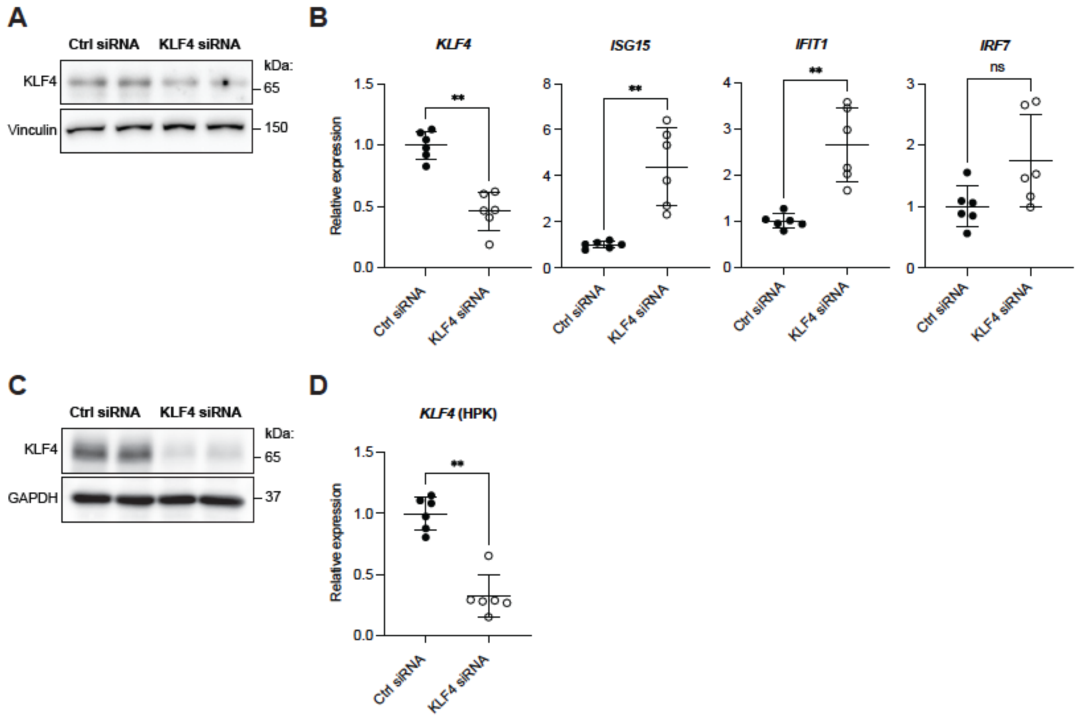


**Supplementary Fig. 6: siRNA-mediated KLF4 knock-down promotes ISG expression**

**A)** Western blot of lysates of HaCaT keratinocytes, transfected with scrambled (scr) or KLF4 siRNA.

**B)** RT-qPCR for *KLF4*, *ISG15*, *IFIT1*, *IRF7* relative to *RPL27* using RNA from HaCaT keratinocytes, transfected with scr or KLF4 siRNA (N = 6).

**C)** Western blot of lysates of HPKs, transfected with scr or KLF4 siRNA.

**D)** RT-qPCR for *KLF4* relative to *RPL27* using RNA from HPKs, transfected with scr or KLF4 siRNA (N = 6; HPKs from two donors).

Data information: Graphs show mean and SD. Non-significant (ns); **P < 0.01 (Mann-Whitney U test (B, D)).

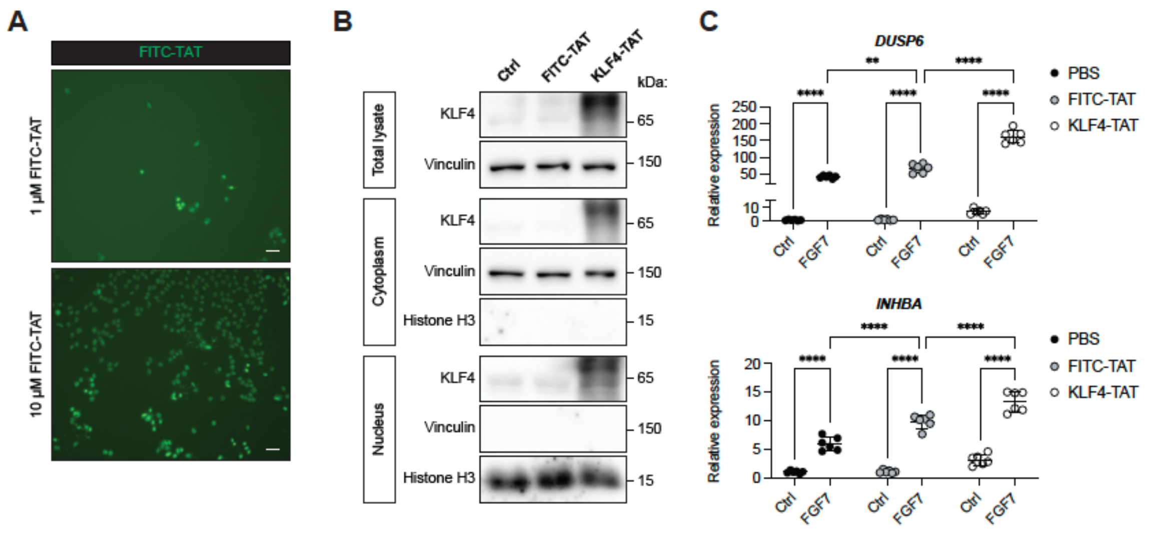


**Supplementary Fig. 7: Delivery of KLF4-TAT into HaCaT keratinocytes promotes expression of *DUSP6* and *INHBA***

**A)** Representative fluorescence images of confluent HaCaT keratinocytes incubated for 2 h with membrane-permeable FITC-TAT (1 or 10 µM). Scale bar: 50 µm.

**B)** Western blot of total, cytoplasmic and nuclear lysates of serum-starved HaCaT keratinocytes, pre-incubated for 2 h with membrane-permeable KLF4-TAT, FITC-TAT or vehicle. Membranes were probed with antibodies against KLF4, vinculin (cytoplasmic marker) and histone H3 (nuclear marker).

**C)** RT-qPCR for *DUSP6* and *INHBA* relative to *RPL27* using RNA from serum-starved HaCaT keratinocytes, pre-incubated with membrane-permeable KLF4-TAT, FITC-TAT or vehicle and treated with FGF7 or vehicle for 6 h (N = 6 per treatment group).

Data information: Graphs show mean and SD. **P < 0.01; ****P < 0.0001 (2-way ANOVA with Bonferroni`s multiple comparisons test).

| gene_id | gene_name | log2 Ratio | pValue | fdr |
|---|---|---|---|---|
| ENSG00000189143 | *CLDN4* | 0.893388898 | 5.32865E-18 | 9.94173E-17 |
| ENSG00000181885 | *CLDN7* | 0.492817055 | 1.35176E-14 | 1.89828E-13 |
| ENSG00000001084 | *GCLC* | 0.459936136 | 8.36012E-10 | 7.20681E-09 |
| ENSG00000023909 | *GCLM* | 0.428416087 | 1.59178E-05 | 7.36142E-05 |
| ENSG00000181019 | *NQO1* | 0.410400345 | 9.99725E-07 | 5.68989E-06 |
| ENSG00000123131 | *PRDX4* | 0.405176346 | 0.000301822 | 0.001088895 |
| ENSG00000157870 | *PRXL2B* | 0.404988143 | 6.60529E-05 | 0.000272991 |
| ENSG00000165672 | *PRDX3* | 0.388067122 | 3.13351E-06 | 1.65482E-05 |
| ENSG00000126432 | *PRDX5* | 0.237860055 | 7.7616E-05 | 0.000316968 |
| ENSG00000156284 | *CLDN8* | 0.231902612 | 0.008911542 | 0.022651046 |
| ENSG00000167815 | *PRDX2* | 0.202505833 | 0.004062385 | 0.011385138 |
| ENSG00000116044 | *NFE2L2* | 0.168533899 | 0.00766578 | 0.019868046 |
| ENSG00000117592 | *PRDX6* | 0.165711257 | 0.0070735 | 0.018509298 |
| ENSG00000117450 | *PRDX1* | 0.126052159 | 0.011176143 | 0.027644021 |

**Supplementary Table 1**

FGF7-induced genes encoding tight junction proteins and genes involved in ROS detoxication.
